# HflX controls hypoxia-induced non-replicating persistence in slow growing mycobacteria

**DOI:** 10.1101/2020.03.13.990168

**Authors:** Jie Yin Grace Ngan, Swathi Pasunooti, Wilford Tse, Wei Meng, So Fong Cam Ngan, Sze Wai Ng, Muhammad Taufiq Jaafar, Huan Jia, Su Lei Sharol Cho, Jieling Lim, Hui Qi Vanessa Koh, Noradibah Abdulghani, Kevin Pethe, Siu Kwan Sze, Julien Lescar, Sylvie Alonso

## Abstract

GTPase HflX is highly conserved in prokaryotes and is a ribosome splitting factor during heat shock in *E. coli.* Here we report that HflX produced by slow growing *M. tuberculosis* and *M. bovis* BCG is a GTPase that plays a critical role in the pathogen’s transition to a non-replicating, drug-tolerant state in response to hypoxia. Indeed, HflX-deficient *M. bovis* BCG (KO) replicated markedly faster in the microaerophilic phase of a hypoxia model, that precipitated entry into dormancy. The KO displayed the hallmarks of dormant mycobacteria including phenotypic drug resistance, altered morphology, low intracellular ATP and up-regulated dormancy *dos* regulon. KO-infected mice displayed increased bacterial burden during the chronic phase of infection, consistent with the higher replication rate observed *in vitro* in microaerophilic phase. Unlike fast-growing mycobacteria, BCG HlfX was not involved in antibiotic resistance under normoxia. Proteomics, pull-down and ribo-sequencing supported that mycobacterial HflX is a ribosome binding protein that controls the translational activity of the cell. Collectively, our study provides further insights into the mechanisms deployed by mycobacteria to adapt to their hypoxic microenvironment.

## Introduction

GTP binding proteins are found across all living kingdoms and are involved in the regulation of many cellular processes. Among which, the small GTPases act as molecular switches that are active or “on” when binding GTP and inactive or “off” when binding GDP. Universally conserved prokaryotic GTPases (ucpGTPases) are a core group of GTPases which are conserved in most prokaryotes, hinting at a critical function in biology (Verstraeten, 2011), though the actual physiological role of most of them has remained elusive. ucpGTPases are characterized by the presence of highly conserved motifs or domains, including the phosphate-loop (P-loop) within the G-domain, a characteristic site where GTP binding and hydrolysis occur (Bourne, 1990; Sprang, 1997).

<u>H</u>igh frequency of lysogenization <u>X</u> (HflX) protein belongs to the superfamily of the Obg-HflX-like ucpGTPases, class of <u>Tra</u>nslation <u>Fac</u>tors (TRAFAC), which have been described to participate in protein translation, likely by providing energy for protein synthesis, or by facilitating the recycling of factors involved in translation (Laalami, 1996; Verstraeten, 2011). HflX was first reported as part of the *hflA* locus in *Escherichia coli* that controls the phage lysis−lysogeny decision process and was thus initially thought to play a role in transposition (Dutta, 2009). However, more recent work has shown that *E. coli* HflX acts as a ribosome-splitting factor under heat shock stress, whereby it binds to and splits stalled 50s ribosomal subunits (Coatham, 2016; Zhang, 2015). In *Staphylococcus aureus*, HflX binds to and dissociates hibernating 100S ribosomes (homodimeric 70S) into 50S and 30S subunits, thereby recycling the pool of ribosomes for new rounds of translation during the stationary phase (Basu, 2017). A recent study has reported a similar ability for HflX expressed by *Mycobacterium abscessus* and *M. smegmatis* to bind to and split ribosomal subunits (Rudra, 2020).

In *Mycobacterium tuberculosis* (Mtb), responsible for human tuberculosis, *hflX* has been categorized as a non-essential gene using a Transposon transposon site hybridization (TraSH) library (Sassetti, 2003). Other studies found that *hflX* gene expression was influenced by a variety of stressors ranging from antibiotics and chemical exposure, to environmental stresses such as nutrient starvation (Andreu, 2008; Boshoff, 2004; Dutta, 2010; FU, 2009; Manjunatha, 2009; Morris, 2005; Sherrid, 2010). Consistently, we previously reported that *hflX* is over-expressed in Mtb exposed to the hostile lysosomal environment of macrophages (Lin, 2016). Transcription of *hflX* was found to be linked with *whiB7*, a transcriptional regulator that controls intrinsic antibiotic resistance and redox homeostasis in Mtb (Burian, 2012; Morris, 2005). Exposure to antibiotics targeting the ribosomal complex including streptomycin, erythromycin, tetracycline, and pristinamycin was found to induce both *whiB7* and *hflX* expression (Hartkoorn, 2012). Consistently, absence of HflX in fast growing mycobacteria species *M. abscessus* and *M. smegmatis* increased antibiotic resistance to macrolide-lincosamide antibiotics (Rudra, 2020).

In this study, we investigated the physiological role of HflX in tubercle bacilli *M. bovis* BCG and *M. tuberculosis*. We report that Mtb/BCG HflX is a GTPase that is involved in response to hypoxia-induced persistence, a non-replicating state that allows tubercle bacilli to persist inside their host for extended periods and become phenotypically antibiotic-resistant (Gold, 2017; Kester, 2014). We provide experimental evidence that HflX interacts with ribosomal subunits and plays a master regulatory role in protein translation during transition to hypoxia.

## Results

### Mycobacterial HlfX is a GTPase with minimal ATPase activity

In the prokaryotic kingdom, HflX is widely distributed and conserved across species (Leipe, 2002; Verstraeten, 2011).The amino acid sequence of *M. tuberculosis* (Mtb) HflX is 100% and 84.5% identical to *M. bovis* BCG HflX and *M. leprae* HflX, respectively, while it shares about 45% identity within the GTPase catalytic site of *E. coli* HflX, including the P loop, Switch I-II, and G1-G5 domains (*Appendix*, Fig. S1A). A three-dimensional computational model was constructed by adopting a previously described strategy (Fischer, 2012). The Phyre2.0 modeling platform was employed to compare the predicted structure of Mtb HflX to that of *E. coli* HflX, whose crystal structure is available (PDB entry: 5ADY) (Zhang, 2015). As expected from the high level of amino-acid identity, a high degree of homology was observed visually and from the low root-mean-square-deviation (RMSD) score, which supports the conservation of HflX structure and function between these two evolutionarily distant prokaryotes (*Appendix*, Fig. S1B).

To investigate whether mycobacterial HflX is a GTPase, codon-optimized BCG/Mtb HflX was expressed in and purified from, *E. coli* (*Appendix*, Fig. S2A&B). Significant GTPase activity in the presence of MgCl_2_ but limited ATPase activity could be observed (Fig. 1A&B; *Appendix*, Fig. S2C). A mutant harboring a triple amino acid substitution (AAY) in the predicted GTPase catalytic site was also generated (*Appendix*, Fig. S2A) and was found to be unable to hydrolyze GTP (Fig. 1A). Furthermore, direct interaction between mycobacterial HflX and GTP hydrolysis product GDP, was demonstrated by isothermal titration calorimetry (ITC) with a dissociation constant Kd at 1.89 µM and a 1:1 stoichiometry (Fig. 1C&D). On the other hand, no significant interaction between the triple mutant HflX and GDP was observed. These data thus establish that HflX produced by Mtb and *M. bovis* BCG is a GTPase with minimal ATPase activity.

**Figure 1.**
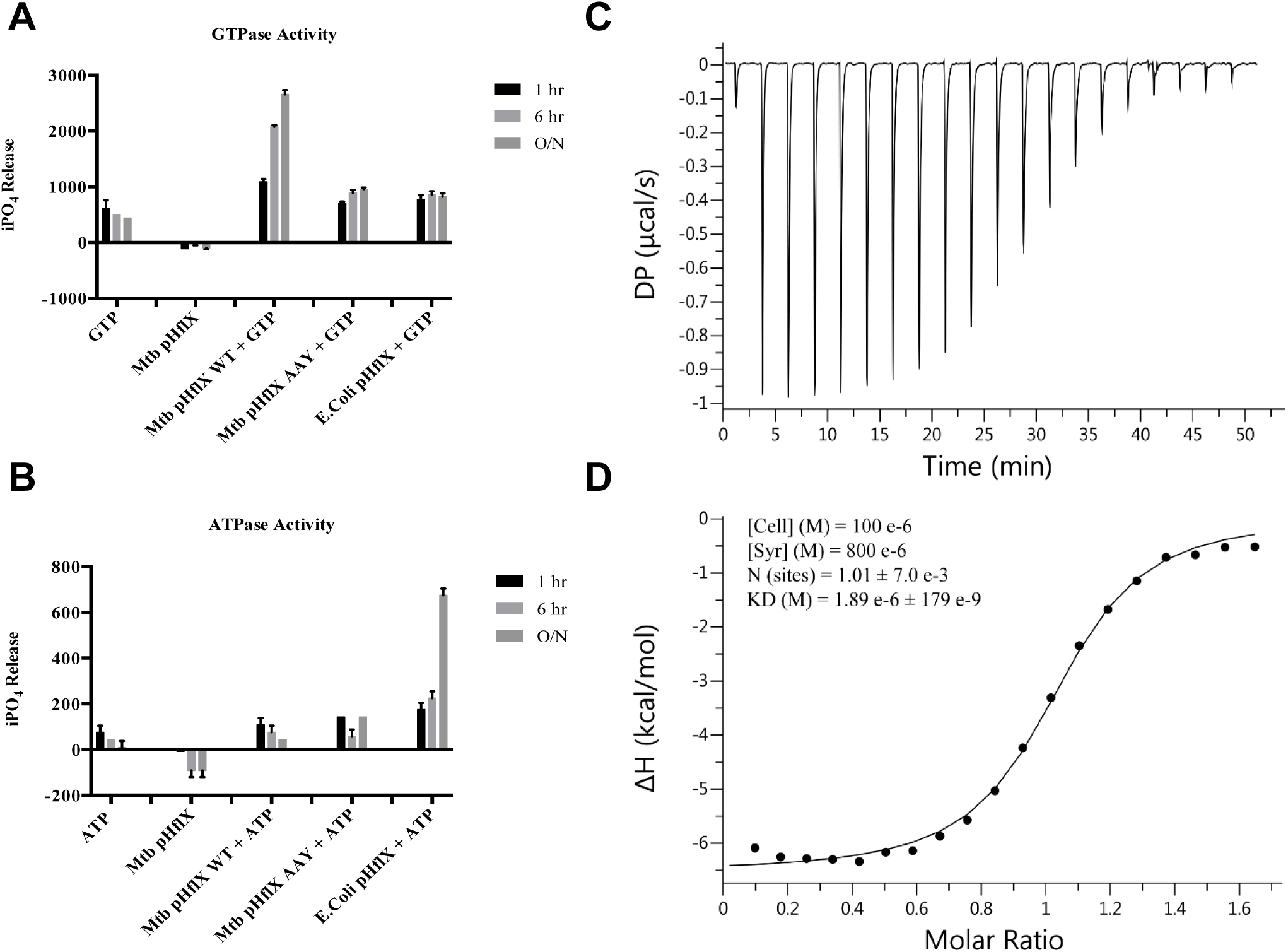
GTPase and ATPase activities of purified mycobacterial HflX. A. Quantification of inorganic phosphate (IPO_4_) released over time in the presence of GTP with Mtb HflX, GTPase-abrogated Mtb HflX AAY or *E.coli* HflX. Data show mean ± SD of three independent experiments. B. Quantification of inorganic phosphate (IPO_4_) released over time in the presence of ATP with Mtb HflX, GTPase truncated Mtb HflX AAY or *E.coli* HflX. Data show mean ± SD of three independent experiments. C, D. Binding of HflX to GDP. (C) Representative differential power (DP) trace of the isothermal titration calorimetry (ITC) experiment. (D) Binding curve of the same experiment, obtained by integrating the DP signal. Two independent experiments were performed. Results from one representative of two independent experiments are shown.

### Mycobacterial HflX is involved in adaptation to hypoxia

*E. coli* HflX has been reported to be a ribosome splitting factor responding to heat shock (Dey, 2018; Zhang, 2015). We thus investigated whether HflX from tubercle bacilli would have a similar function. A *M. bovis* BCG HflX null mutant (Δ*hflX*) and its complemented strain (Δ*hflX::phflX*) were constructed (*Appendix*, Fig. S3A). RT-PCR revealed undetectable levels of *hflX* mRNA in BCG Δ*hflX*, while the complemented strain displayed *hflx* mRNA level similar to the parental strain (*Appendix*, Fig. S3B). Comparable growth kinetics were observed amongst WT, Δ*hflX*, and Δ*hflX::phflX* strains when cultured in standard 7H9 liquid culture medium (*Appendix*, Fig. S3C), supporting that HflX is non-essential for *in vitro* growth in rich aerated (normoxic) culture medium at 37°C.

Mycobacterial HflX was not found to play a role during heat shock as evidenced by comparable number of colony-forming units (CFU) amongst the WT, Δ*hflX*, and complemented strains (Fig. 2A). Furthermore, codon-optimized BCG HflX or homologous *E. coli* HflX were expressed in Δ*hflx E. coli* under the control of an arabinose-inducible promoter (*E. coli ΔhflX::pBCGhflX and E. coli ΔhflX::hflX*). The *hflX* mRNA levels in both strains were comparable and about 100 times higher than the endogenous level measured in WT *E. coli* (*Appendix*, Fig. S3D). Upon heat shock, expression of homologous HflX partially restored parental survival, while codon-optimized BCG HflX did not confer protection to the Δ*hflX E. coli* strain (Fig. 2B). Thus together, these observations support that mycobacterial HflX is unlikely to be involved in the heat shock response.

**Figure 2.**
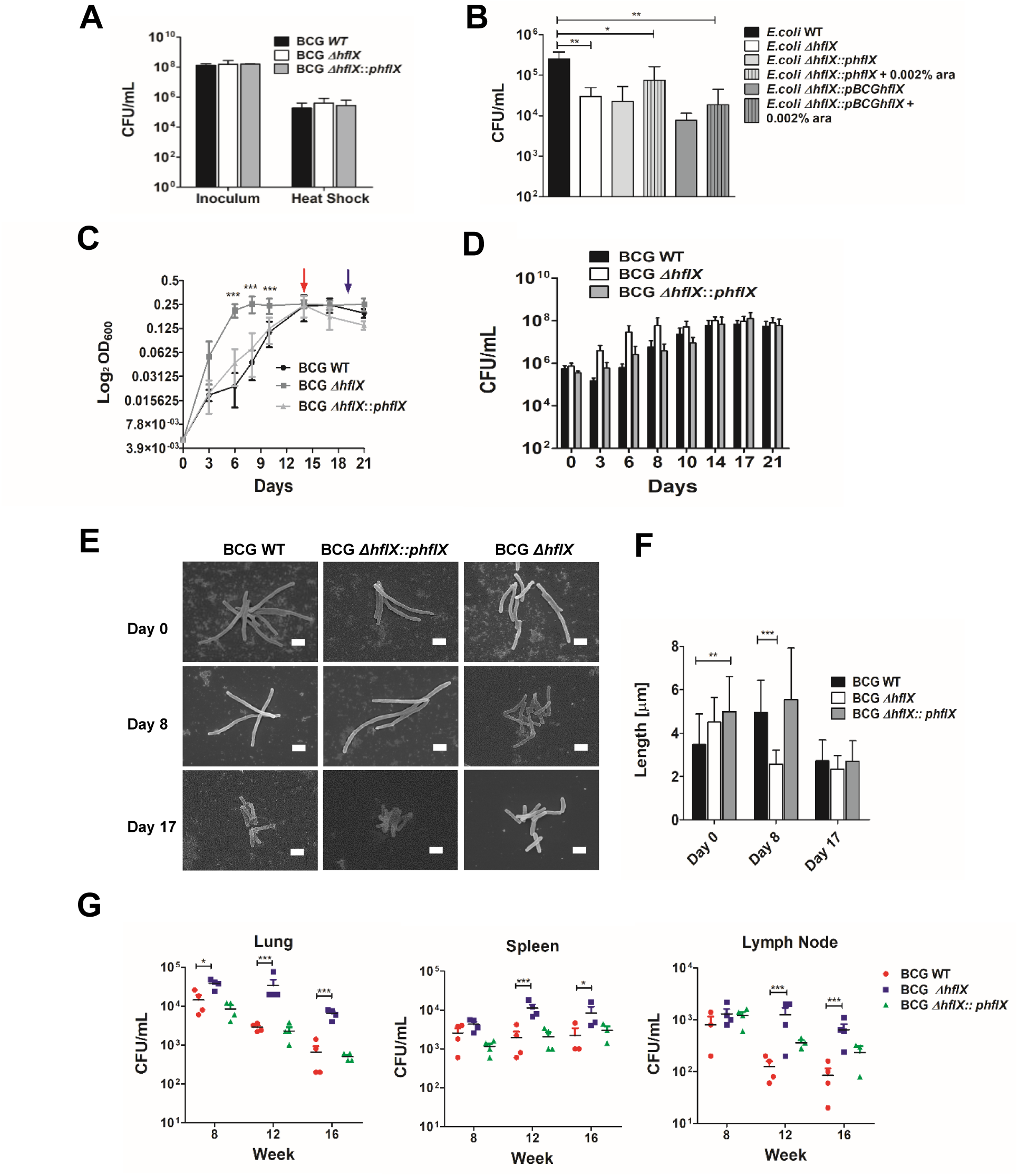
Infection profile of BCG ΔHflX in the mouse model. A. CFU counts from BCG WT, Δ*hflX* and complemented strains before and after heat shock stress as described in methods. Data show mean ± SD of three independent experiments. B. CFU counts from *E.coli* WT, Δ*hflX*, and Δ*hflX* complemented with homologous HflX or with BCG codon-optimized HflX after heat shock stress as described in methods. Data show mean ± SD of three independent experiments. C. OD_600_ of BCG WT, Δ*hflX* and complemented strains in the gradual hypoxia Wayne model. Red arrow, WT reached NRP-1; Blue arrow, WT reached NRP-2. Data show mean ± SD of four independent experiments. D. CFU from BCG WT, Δ*hflX* and complemented strains in the gradual hypoxia Wayne model. Data show mean ± SD of four independent experiments. E. Representative images of BCG WT, Δ*hflX* and complemented strains obtained by scanning electron microscopy on day 0, 8 and 17 of the Wayne model. Scale bar = 1µm. F. Average length of BCG WT, Δ*hflX* and complemented strains based on 20 bacteria counted for each strain and on day 0, 8 and 17 of the Wayne model. G. CFU counts from C57BL/6 mice infected with BCG WT, Δ*hflX* and complemented strains. Organs were harvested at week 8, 12 and 16 post-infection. One representative experiment out of two is shown. Data information: All data show mean ± SD *P < 0.05, **P < 0.01, ***P < 0.001. Panels (B, F, G) one-way ANOVA with Bonferroni post-test.

To probe a possible physiological role of mycobacterial HflX during adaptation to other stresses, the BCG WT, Δ*hflX*, and complemented strains were grown under various conditions, including macrophage infection, nutrient starvation (Loebel *in vitro* model), and gradual oxygen depletion (Wayne *in vitro* model). No significant difference among the three strains was observed during macrophage infection and under nutrient starvation (*Appendix*, Fig. S4A&B). Growth profiles were then monitored in the gradual oxygen depletion model (aka Wayne model). In this *in vitro* model, mycobacteria growth is characterized by two stages, namely the non-replicating persistence stage 1 (NRP-1) or microaerophilic stage, during which mycobacteria slow-down replication while oxygen gets progressively depleted in the sealed tube (<1.0% O_2_); and the non-replicating persistence stage 2 (NRP-2), characterized by an oxygen tension below 0.06% and where mycobacteria have stopped replicating and enter a dormant state (Wayne, 1996; Wayne, 2001). In this model, during the NRP-1 stage (days 3-8), the BCG Δ*hflX* culture was found to grow faster than the WT and complemented strains, as evidenced by significantly higher OD_600nm_ values and CFU counts (Fig. 2C&D). The growth of BCG Δ*hflX* ceased on day 8 onwards as evidence by plateaued OD_600nm_ values, and reached the NRP-2 stage at day 14 (as indicated by complete decolorization of methylene blue indicator), while the WT and complemented strains reached NRP-2 by day 18 and 17, respectively (Fig. 2C&D). The differential growth kinetic profile observed in the Wayne model with the HflX-deficient strain thus pointed at a role for HflX in controlling growth rate during the microaerophilic phase and entry of mycobacteria into the non-replicating state.

Changes in cell morphology have been reported previously for non-replicating mycobacteria grown under hypoxic conditions and external acidification, including thickening of the cell wall, size reduction, and ovoid cell formation (Jakkala, 2019; Shleeva, 2011; Velayati, 2011). The size and morphology of BCG Δ*hflX* mycobacteria grown in the Wayne model were studied by scanning electron microscopy. A 43% reduction in size was observed with BCG Δ*hflX* at day 8 (NRP-1) compared to its size at day 0, while the average size of WT and complemented strains was comparable to day 0 (Fig. 2E,F). At day 21 (NRP-2), the size of WT and complemented strains decreased significantly compared to the NRP-1 stage and day 0, reaching a size that was similar to that measured with Δ*hflX* strain (Fig. 2E,F). These observations, therefore, suggested that the Δ*hflX* mutant displayed a non-replicative phenotype earlier than the WT and complemented strains.

We were next interested to test whether HflX impacts mycobacterial growth in a mammalian host where oxygen saturation ranges between 1-14% depending on the organ, thereby likely exposing mycobacteria to microaerophilic environments (Carreau, 2011; Liu, 2011). Upon intratracheal infection, the number of CFUs recovered at weeks 2 and 4 from the lungs, spleen and lymph nodes of mice infected with WT, Δ*hflX* and complemented strains were mostly comparable (*Appendix* Fig. S4E). In contrast, from week 8 onwards, which corresponds to the chronic phase of infection triggered by the host adaptive immunity (Köhler, 1975; Nicolle, 2004), the bacterial loads measured in these organs were consistently higher in mice infected with Δ*hflX* compared to the parental and complemented strains (Fig 2G). These findings were consistent with the higher replication rate observed with Δ*hflX* mutant during microaerophilic phase of the *in vitro* hypoxic Wayne model, and supported a role for HflX in the physiological response of mycobacteria to low oxygen tension environments that are encountered during the course of infection in the human host.

### Absence of HflX impairs the energetic status of hypoxic mycobacteria

We previously reported that under gradual oxygen depletion, the intracellular ATP level drops significantly in mycobacteria (Rao, 2007). We thus monitored the intracellular ATP levels in BCG Δ*hflX* grown in the Wayne model. While comparable intracellular ATP levels were measured at day 0 for the WT, Δ*hflX*, and complemented strains, significantly lower ATP levels were obtained with the Δ*hflX* strain at all the subsequent time points (Fig. 3A). Furthermore, we also determined the membrane potential (Δψ) of the three strains using cationic fluorescent dye DiOC_2_, as previously described (Rao, 2007; Vaara, 1992). Negative controls consisted of cultures incubated with proton-ionophore cyanide m-chlorophenyl hydrazine (CCCP) that dissipates the transmembrane proton gradient (ΔpH) component of proton motive force (PMF). BCG Δ*hflX* displayed an overall 17-46 % increase of Δψ values compared to WT and complemented strains, with a 2-fold increase on day 3 (Fig. 3B), suggesting that the plasma membrane of BCG Δ*hflX* is hyperpolarized during growth in the Wayne model. Of note, the addition of CCCP only caused a slight reduction of RFU, presumably reflecting an incomplete depolarized state of the membrane.

**Figure 3.**
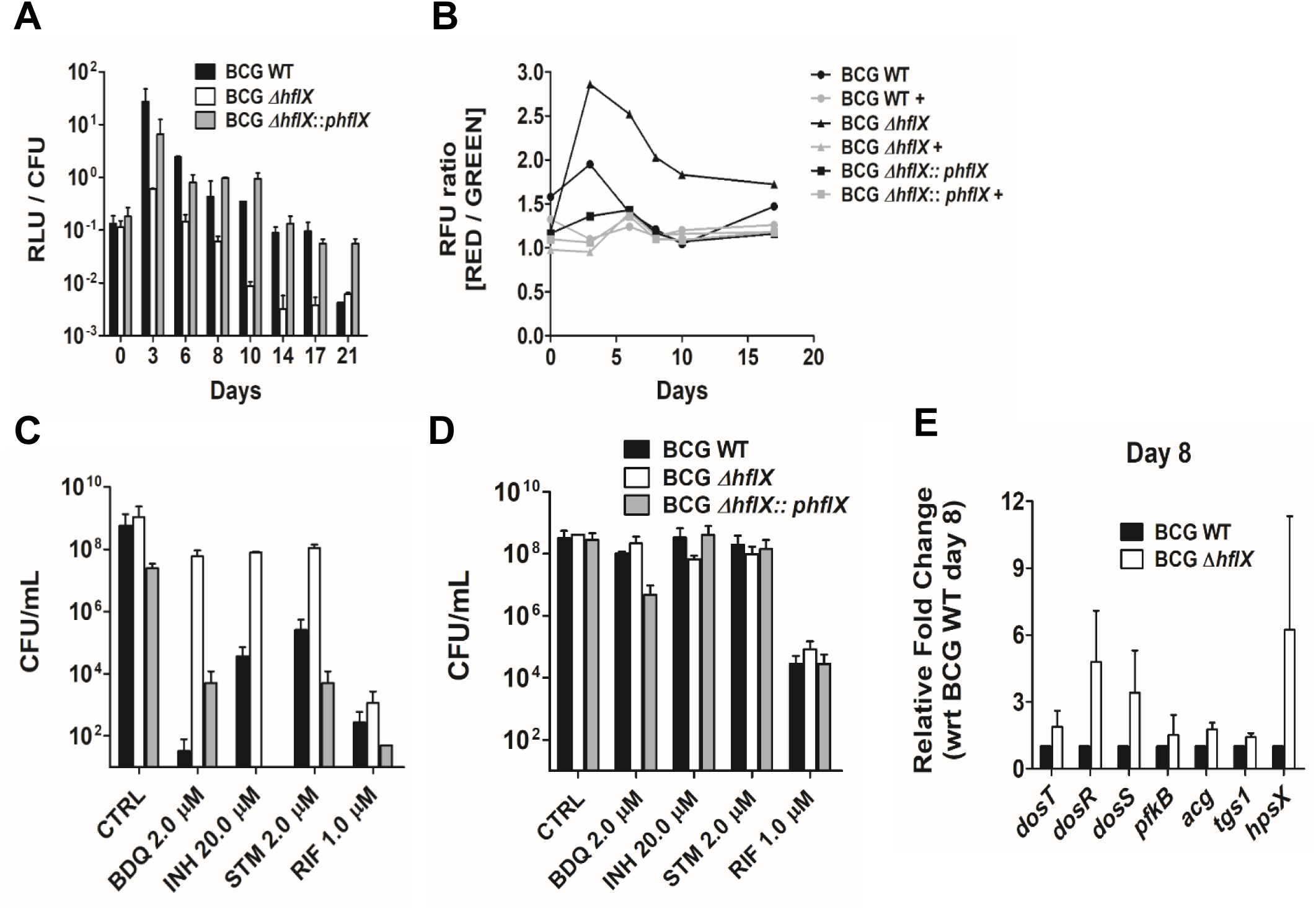
Energetic status, drug susceptibility and expression of the *dos* regulon in BCG □HflX. A. Intracellular ATP level in BCG WT, Δ*hflX* and complemented strains grown in the Wayne model. Data show mean ± SD of three independent experiments. B. Membrane potential of BCG WT, Δ*hflX* and complemented strains grown in the Wayne model. + CCCP as a membrane disruptor positive control. One representative of two independent experiments shown. C. CFU from BCG WT, Δ*hflX* and complemented strains on day 8 of the Wayne model treated with various drugs. Data show mean ± SD of two independent experiments. D. CFU from BCG WT, Δ*hflX* and complemented strains on day 17 of the Wayne model treated with various drugs. Data show mean ± SD of two independent experiments. E. Relative gene expression of a subset of *dos* regulon genes measured by RT-PCR in BCG Δ*hflX* compared to WT on day 8 of the Wayne model. Data show mean ± SD of two independent experiments.

Altogether, our data indicated that under gradual oxygen depletion, BCG Δ*hflX* displays significantly lower ATP levels compared to WT and complemented strains that could explain its earlier entry into a non-replicating state. The membrane hyperpolarization observed may reflect a compensatory mechanism to maintain the PMF for *de novo* ATP synthesis critical for mycobacterial survival (Srinivasa PS Rao et al., 2007).

### BCG Δ*hflX* exhibits phenotypic drug resistance in NRP-1 that correlated with up-regulation of the *dos* regulon

The phenotypes displayed by Δ*hflX* in Wayne model prompted us to investigate the drug susceptibility profile of this mutant. Phenotypic drug resistance of non-replicating mycobacteria induced under hypoxia, nutrient starvation, or stationary phase has indeed been well described (Franzblau, 2012; Lakshminarayana, 2015; Rao, 2007). This phenomenon is believed to explain the prolonged chemotherapy necessary to achieve sterility and cure in TB patients (Davies, 2010; McCune, 1966; McCune, 1956; Nuermberger, 2004). Here, we investigated the susceptibility of BCG Δ*hflX* to various anti-mycobacterial drugs with different mechanisms of action, namely bedaquiline BDQ, isoniazid INH, streptomycin STM, rifampicin RIF, chloramphenicol CM, and ethambutol ETB. WT, Δ*hflX* and complemented strains grown under aerobic conditions displayed comparable minimum inhibitory concentrations (MIC) (*Appendix*, Table S1). In NRP-1 of the Wayne model, however, BCG Δ*hflX* displayed high resistance to BDQ, INH, and STM, with only 1-2 log_10_ decrease in CFU/mL compared to the drug-free control, while WT and complemented strains displayed between 3-8 log_10_ reductions in CFU/mL when exposed to these drugs (Fig. 3C). Of note, BCG Δ*hflX* did not display increased drug resistance to RIF compared to WT and complemented strains. In the NRP-2 stage, all three strains were resistant to BDQ, INH, and STM and remained susceptible to RIF (Fig. 3D), consistent with previous studies reporting that RIF is very effective at killing non-replicating mycobacteria (Iacobino, 2017; Iacobino, 2016; Tomasz, 1970). The lack of killing efficacy of non-replicating mycobacteria observed with BDQ, previously shown to kill both actively replicating and non-replicating mycobacteria (Andries, 2005; Haagsma, 2011), may be explained by the fact that this drug exerts a delayed killing (Koul, 2014), and that 5-day incubation may not be sufficient to observe significant killing of non-replicating mycobacteria (Piccaro, 2015).

Furthermore, the two-component system DosS/T-R has been known to mediate mycobacteria transition to a non-replicating state in response to various stresses, including low oxygen tension (Bretl, 2011; Gautam, 2014; Kendall, 2004; Sharma, 2016). The sensory kinases DosS or DosT activate transcriptional regulator DosR by phosphorylation, leading to the transcription of a 48 gene-regulon (aka *dos* regulon) (Bagchi, 2005; Roberts, 2004; Saini, 2004; Sousa, 2007). We thus examined the transcriptional activity of the *dos* regulon in BCG Δ*hflX*. A significant increase in the transcription level of a number of *dos* regulon genes (*dosR*, *dosS*, *dosT*, and *hspX*) was found with the Δ*hflX* mutant in the NRP-1 phase (day 8) of the Wayne model with average fold-increases of 5X, 3X, 1.5X and 6X respectively compared to WT (Fig. 3E and *Appendix* Fig. S4 C&D).

Together, both the phenotypic drug resistance profile and up-regulation of the *dos* regulon in the NRP-1 stage of the Wayne model, further supported that in this gradual oxygen depletion model, BCG Δ*hflX* enters a non-replicating state earlier than its parental counterpart.

### Differential proteomic profile in BCG Δ*hflX* in response to hypoxia

As a ribosome-splitting factor, *E. coli* HflX influences the translational activity in this bacterium. To understand the mechanisms by which HflX plays a role in triggering the non-replicating state in tubercle bacilli, we employed tandem mass tag mass spectrometry (TMT-MS) to analyze the relative protein abundance in Δ*hflX* BCG compared to the parental strain when grown under hypoxic conditions. Results indicated differential protein content between WT and Δ*hflX* at all the time points tested (day 0, day8, and day 17) (Fig. 4; *Appendix*; Tables S2-S4). At day 0, 66 and 56 proteins were down- and up-regulated in BCG Δ*hflX*, respectively (Fig. 4A). According to gene ontology analysis, significant enrichment was found for down-regulated proteins involved in lipid biosynthesis and metabolism, cell wall components biogenesis and assembly, leucine biosynthesis, protein folding and response to copper ion (Fig. 4D). At day 8, extensive differential protein abundance was seen between the BCG Δ*hflX* and WT, with 151 up-regulated and 201 down-regulated proteins (Fig. 4B). Among the 151 up-regulated proteins, 15 were encoded by genes from the *dos* regulon (*Appendix*, Table S3, bolded), in line with the observed up-regulation of *dos* regulon genes at the transcriptional level. Furthermore, and consistent with a potential regulatory role of HflX in translational activity of the bacterium, the day-8 Δ*hflX* sample was enriched in ribosomal subunits (*Appendix*, Table S3, highlighted in yellow) and in proteins involved in formation of the hibernating ribosome (HPF, RafH) (*Appendix*, Table S3, highlighted in green). Also, in that sample, many of the up-regulated proteins were assigned to central metabolism (PfkB, Gap, FabG1, CitA, AceA, Icd2, and SdhB) (*Appendix*, Table S3). The proteins that were down-modulated at day 8 were similar to those down-regulated at day 0 (Fig. 4E). Interestingly, at day 0 and day 8, BCG Δ*hflX* displayed significantly lower amounts of seven proteins (Mas, FadD26, FadD29, FadD22, PpsB, PpsC, PpsD) (*Appendix*, Table S2) encoded by the operon involved in the synthesis of phenolphthiocerol and phthiocerol dimycocerosates (PDIM), a major long-chain fatty acid component of the cell wall in mycobacteria (Azad, 1997; Jackson, 2014; Jackson, 2007). Finally, at day 17, where both strains have reached a non-replicating state (NRP-2), 15 proteins involved in the response to starvation and copper ion were found to be up-regulated in BCG Δ*hflX*, while 27 proteins involved in sulfur compound metabolism, cysteine synthesis, and oxidoreductase activity were down-regulated (Fig. 4C&F, *Appendix*, Table S4).

**Figure 4.**
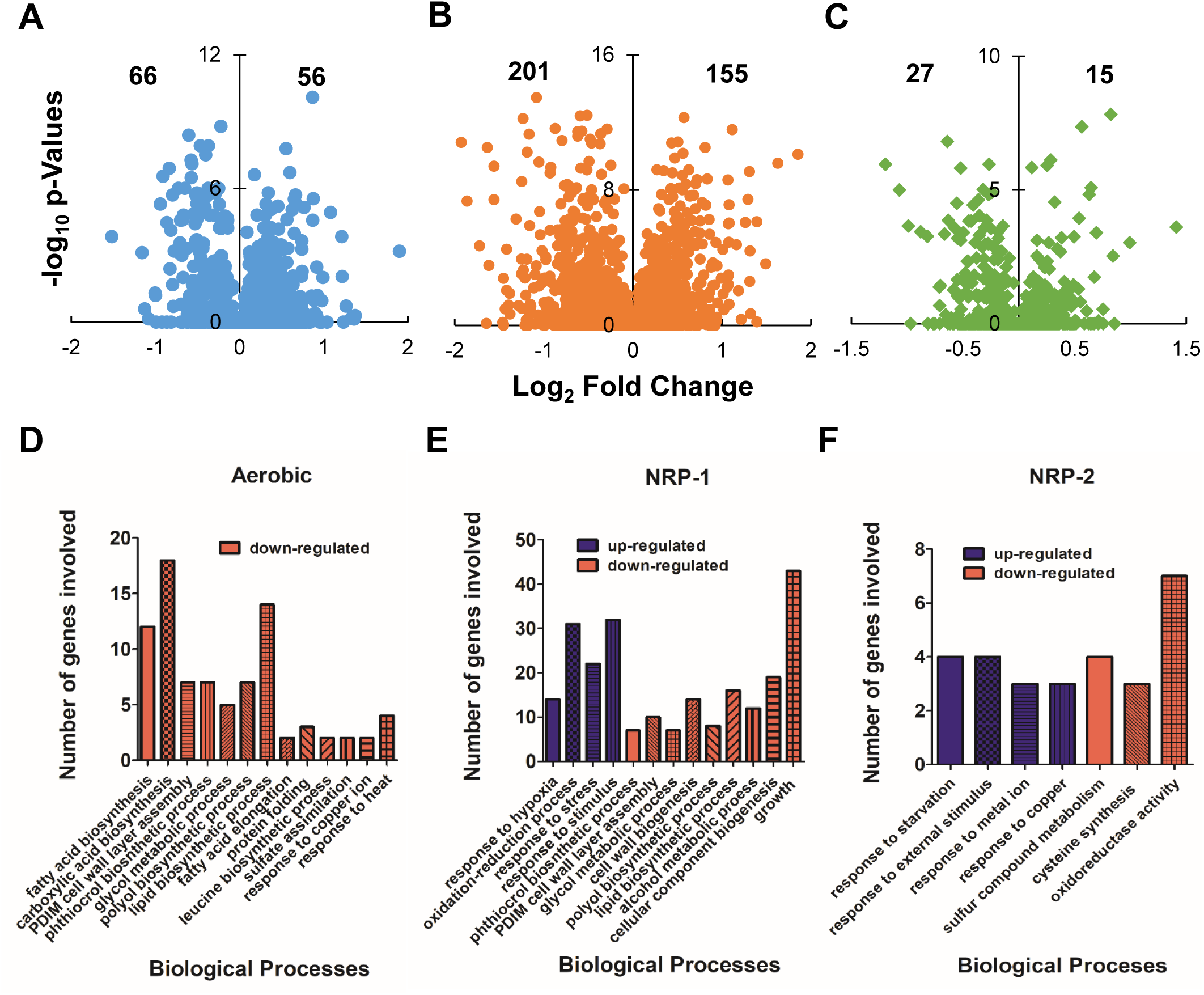
Differential protein expression in BCG Δ*hflX* in response to hypoxia. A-C. Volcano plot of differentially expressed proteins in BCG Δ*hflX* compared to WT on day 0 (A, normoxia), day 8 (B, NRP-1) and day 17 (C, NRP-2) of Wayne model. D-E. Gene ontology analysis (biological functions) of the differentially expressed proteins in BCG Δ*hflX* compared to WT on day 0 (D, normoxia), day 8 (E, NRP-1) and 17 (F, NRP-2) of Wayne model with false discovery rate < 0.05.

Together, this proteomic approach revealed massive changes at the protein level in the BCG Δ*hflX* mutant compared to WT, particularly at day 8 of the Wayne model, which supports a master regulatory role for HflX in response to hypoxia.

### Mycobacterial HflX interacts with ribosomal subunits and regulates protein translation

*E.coli* HflX binds at the E-site of 70S bacterial ribosomes and induces split into 50S/30S ribosomal subunits upon GTP hydrolysis (Coatham, 2016). Using a biochemical approach, a recent study reported that HflX produced by *M. abcessus* and *M. smegmatis* is also a ribosome splitting factor (Rudra, 2020). To investigate whether HflX expressed by slow growing *M. bovis* BCG interacts with ribosomal subunits, we conducted a cell-based pull-down experiment combined with LC/MS analysis using a home-made anti-HflX monoclonal antibody (*Appendix*, Fig S2D&E). Results showed that the pull-down fraction was enriched in ribosomal proteins, namely S6 and S17 of the 30S ribosomal subunits; and L27 and L30 of the 50S ribosomal subunits, thus supporting that HflX binds to ribosomes (Table 1).

**Table 1.**
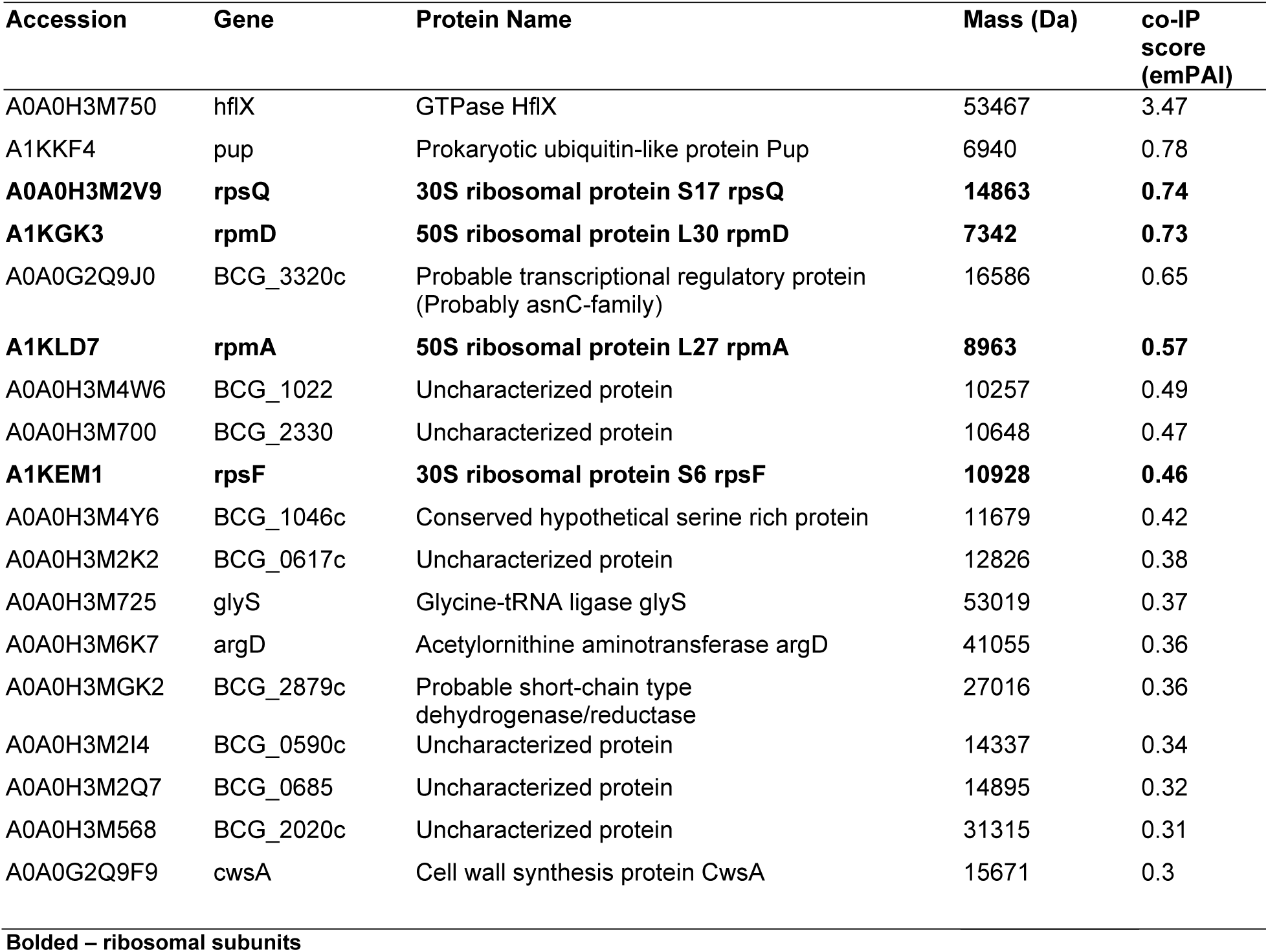
Protein candidates identified from pull-down experiment with anti-HflX antibody.

To further confirm the regulatory role of mycobacterial HflX in protein translation in response to hypoxia, BCG Δ*hflX* or WT bacteria were harvested at day 8 in the Wayne model and were analyzed by ribo-sequencing. This approach revealed the presence of ribosomally protected footprints covering canonical non-coding RNAs (ncRNAs) such as tRNAs and rRNAs, a feature previously reported in a ribo-seq study conducted in *Mycobacterium abcessus* (Miranda-CasoLuengo, 2016). We found an enrichment in tRNA ribosome footprints in the BCG Δ*hflX* compared to WT, while total RNA-seq indicated that the percentages of reads mapped to the respective regions (tRNA, rRNA, CDS and Other) were similar between both strains (Fig.5A, Appendix Fig. S6). Otherwise, the percentage of reads mapped to rRNA, CDS and 5’ and 3’ untranslated regions (Others) by Ribo-seq were generally higher in WT (Fig. 5A). The translation efficiency (TE) of 1,361 coding sequences (CDS) was found to be significantly (Log2TE>1, Log2TE<-1) different between Δ*hflX* and WT, among which 781 had a lower TE in Δ*hflX*, representing approx. 60% of the CDS (Fig. 5B). Interestingly, genes coding for ribosomal subunits were those with the greatest increase in TE in Δ*hflX* (Fig. 5C). This finding was consistent with our proteomics data, and suggested that in the absence of HflX, less free ribosome subunits are available inside the cell, leading bacteria to up-regulate the corresponding genes. Genes involved in metabolic pathways including carbon metabolism, citrate cycle and oxidative phosphorylation were also found to have their translation efficiency up-regulated in Δ*hflX* (Fig 5C). This again may represent a feedback response to the lower ATP pool measured in Δ*hflX*. Among the genes whose translation efficiency was significantly down-regulated in Δ*hflX*, the PPE family genes, genes involved in response to stimuli, and two-component systems were enriched (Fig. 5D).

**Figure 5.**
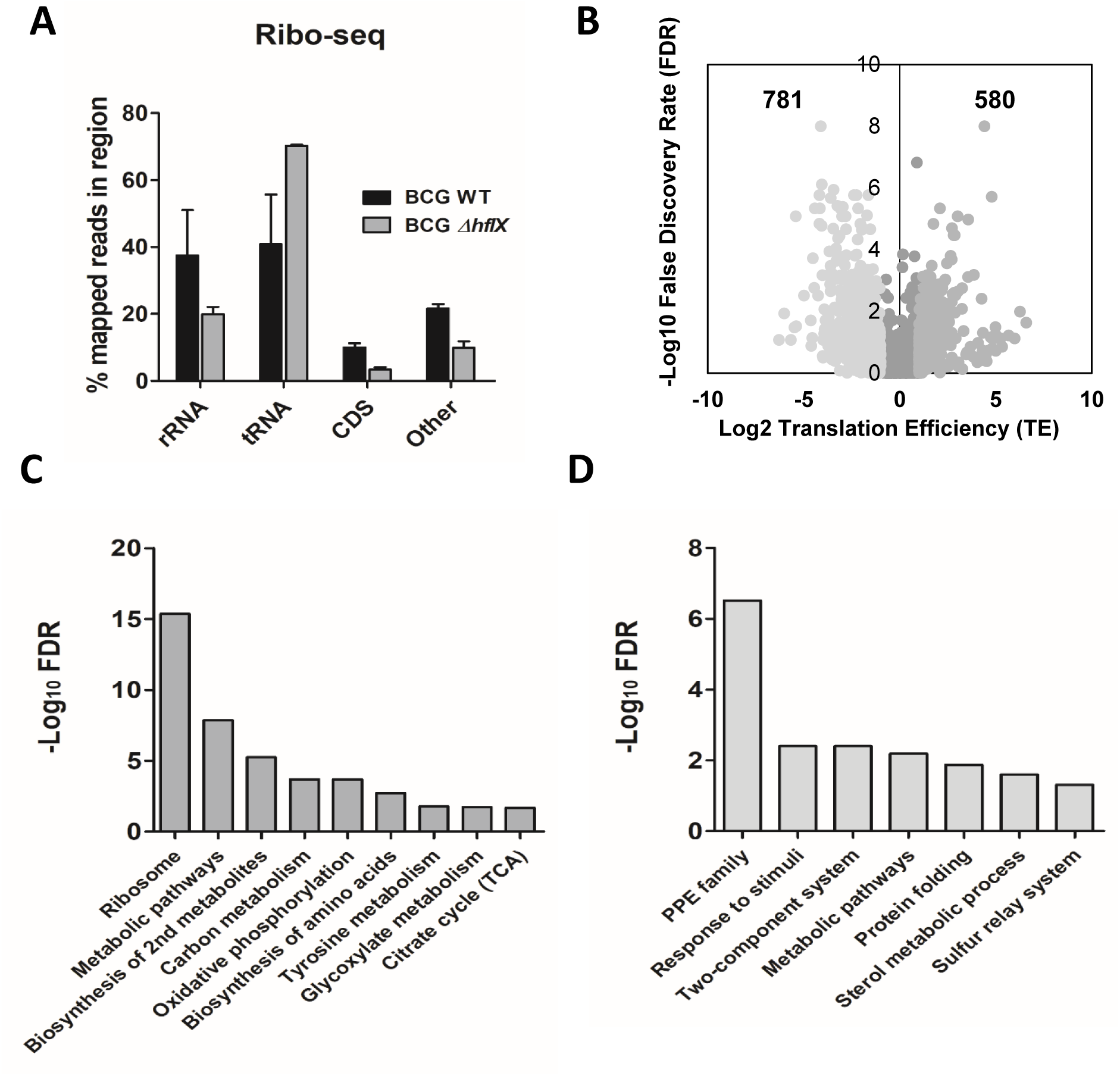
Translational activity in BCG ΔhflX. A. Ribosome sequencing data showing the percentage of mapped reads to respective regions of BCG Δ*hflX* compared to WT on day 8 (NRP-1) of Wayne model. CDS: coding sequence; Other: 5’ and 3’ untranslated regions (UTR). Data show mean ± SD of two independent experiments. B. Volcano plot of differential translation efficiency (TE) between BCG Δ*hflX* and WT on day 8 (NRP-1) of Wayne model. Highlighted in light grey: False discovery rate <0.05. C-D. Gene ontology analysis and KEGG pathways analysis of the differentially translated genes in BCG Δ*hflX* compared to WT on day 8 (NRP-1) of Wayne model. Enriched pathways were selected where false discovery rate was <0.05.

Together, these data support that mycobacterial HflX interacts with ribosomes and plays a regulatory role in protein translation under hypoxic stress.

## Discussion

Mtb can survive for decades in a dormant state, causing a clinically asymptomatic, non-infectious form of the disease that is known as latent TB infection (LTBI) (Dye, 1999; Parrish, 1998). It is estimated that about one-third of the world’s population has LTBI, providing a large reservoir for reactivation to active, contagious disease (WHO., 2019; Veatch and Kaushal., 2018). The ability of dormant Mtb to exhibit a form of non-inheritable resistance to most of the currently available anti-TB drugs (aka phenotypic drug resistance) explains the long treatment regimen needed to achieve sterilization, and has impeded the efforts in TB elimination (Bloom, 1992; Gomez, 2004; Parrish, 1998). During latent infection, non-replicating Mtb bacilli localize within granuloma, an organized structure of immune cells intended to constrain the infection (Dheda, 2005; Orme, 2014; Russell, 2007). The hypoxic microenvironment of granuloma is believed to trigger replication arrest in pathogenic mycobacteria (Dannenberg, 1993; Manabe, 2006; Rustad, 2009; Wayne, 2001). The dormancy survival regulon, aka *dos* regulon, is regulated by the two-component system Dos S/T and DosR and comprises 48 genes, which have been shown to be essential for hypoxic survival (Boon, 2002; Leistikow, 2010; Park, 2003; Roberts, 2004; Sherman, 2001). However, the molecular mechanisms involved in the hypoxic response and replication arrest of pathogenic mycobacteria have remained elusive. Our present work has identified the highly conserved GTPase HflX as a novel mycobacterial factor that plays an important role in the pathogen’s response to its hypoxic environment. Our findings are in line with previous reports on the role of HflX in stress adaptation in other distantly related microorganisms (Basu, 2017; Zhang, 2015). However, instead of heat shock, mycobacterial HflX responded specifically to oxygen limitation, a physiologically relevant stress that mycobacteria encounter in their host environment. Our data support that HflX regulates the translational activity in slow growing pathogenic mycobacteria, and controls entry into the non-replicating state. We further showed that BCG/Mtb HflX is a ribosome-interacting protein, as evidenced by the enrichment in ribosomal subunits in the pull-down fraction (Table 1). This was consistent with the ribosome-splitting activity of HflX described in *E. coli* and *S. aureus* (Basu, 2017; Zhang, 2015), as well as in fast growing mycobacteria species (NTM) *M. abscessus* and *M. smegmatis* (Rudra, 2020). Interestingly, while the latter study reported that HflX-deficient *M. smegmatis* and *M. abscessus* displayed resistance to macrolide-lincosamide, we did not observe any drug resistance phenotype with Δ*hflX* BCG mutant grown in normoxia, including macrolides such as erythromycin (Table S1). Macrolide-lincosamide has been used effectively to treat non-tuberculous mycobacteria (NTM) infections (Binder, 2013; Maxson, 1994; Mushatt, 1995). However, mycobacteria from the Mtb complex (which includes *M. tuberculosis*, *M. bovis* BCG and others, but not *M. smegmatis* or *M. abscessus*) have been found to be intrinsically resistant to macrolides due to the presence of Erm methyltransferase (ErmMT) that confers resistance to macrolide-lincosamide-streptogramin (MLS) by methylation of 23S rRNA (Andini, 2006; Buriánková, 2004). Of note, part of the *ermMT* locus has been deleted in the vaccine strain *M. bovis* BCG Pasteur, making this strain susceptible to erythromycin. Maintenance of erythromycin susceptibility in BCG Δ*hflX* mutant suggests a differential role of HflX between tubercle bacilli and NTM in adaptation to stress.

The hallmark of non-replicating bacilli includes low energy profile and global protein synthesis down-regulation (Hu, 1998; Ignatov, 2015; Schnappinger, 2003; Shi, 2005). In *E. coli*, global translation shut down is associated with ribosome dimerization into a 100S ribosome species, which is translationally inactive when conditions are not favorable for bacterial growth (Gohara, 2018; Starosta, 2014; Wada, 1995; Yamagishi, 1993). Under hypoxic stress, Mtb 70S ribosomes do not dimerize into 100S but associate with hibernating promoting factor (HPF) and ribosome-associated-factor-during-hypoxia (RafH) into a stable complex (Li, 2015; Mishra, 2018; Trauner, 2012). Ribosome stabilization is a strategy deployed by bacteria for stress management, so when cellular conditions become favorable, the hibernating ribosomes get disassembled and quickly recycled for new rounds of translation (El-Sharoud, 2004; Gohara, 2018). The plasticity of hibernating ribosome disassembly has been proposed to play an essential role in the TB disease reactivation process (Sawyer, 2018; Trauner, 2012). It has been shown that *E.coli* and *S. aureus* HflX rescued stalled ribosomes and hibernating 100S ribosomes by splitting them into the 50S and 30S subunits, allowing translation to resume (Basu, 2017; Zhang, 2015). Our proteomics and ribo-seq data indicated an increased abundance in HPF and RafH, and downregulation of 30% of the total translatome, respectively, in BCG Δ*hflX* under oxygen limitation. We thus propose a model whereby under hypoxia, HflX controls the amounts of ribosomal subunits available for translation by splitting hibernating ribosomes and/or stalled ribosomes, thereby controlling the overall translational activity, hence entry into the non-replicating state (Fig. 6). Absence of HflX leads to accumulation of hibernating and stalled ribosomes, precipitating entry into a non-replicating state. The fact that HflX is dispensable in normoxia suggests that either the amount of hibernating ribosomes and stalled ribosomes is negligible, or other splitting factors are at play. The increased amount of individual ribosomal subunits observed in the HflX-deficient mutant could result from a compensatory mechanism that aims to overcome the overall translation shutdown. Alternatively or in addition, Mtb/BCG HflX may also be directly involved in the biogenesis of ribosomal subunits as proposed for several bacterial GTPases, among which many are from the TRAFAC class (Bennison, 2019; Britton, 2009; Campbell, 2008). Furthermore, the extensive changes in proteomic and Ribo-seq profiles observed with BCG Δ*hflX* (including proteins involved in various cellular processes such as central metabolism and cell wall synthesis), coupled with the higher replication rate during the microaerophilic phase and the increased bacterial burden in the chronic phase of infection in mice, suggest a master regulatory role for HflX in mycobacteria’s response to their hypoxic environment. Consistently, some of the TRAFAC-GTPases have been implicated in various cellular processes such as cell wall metabolism, chromosome segregation, and cell division initiation (Britton, 2000; Britton, 1998; Caldon, 2003; Cladière, 2006; Foti, 2007; Gollop, 1991). Whether these pleiotropic effects are a downstream consequence of the regulatory role of HlfX in the bacterium’s translational activity or are independent of it remains to be investigated.

**Figure 6.**
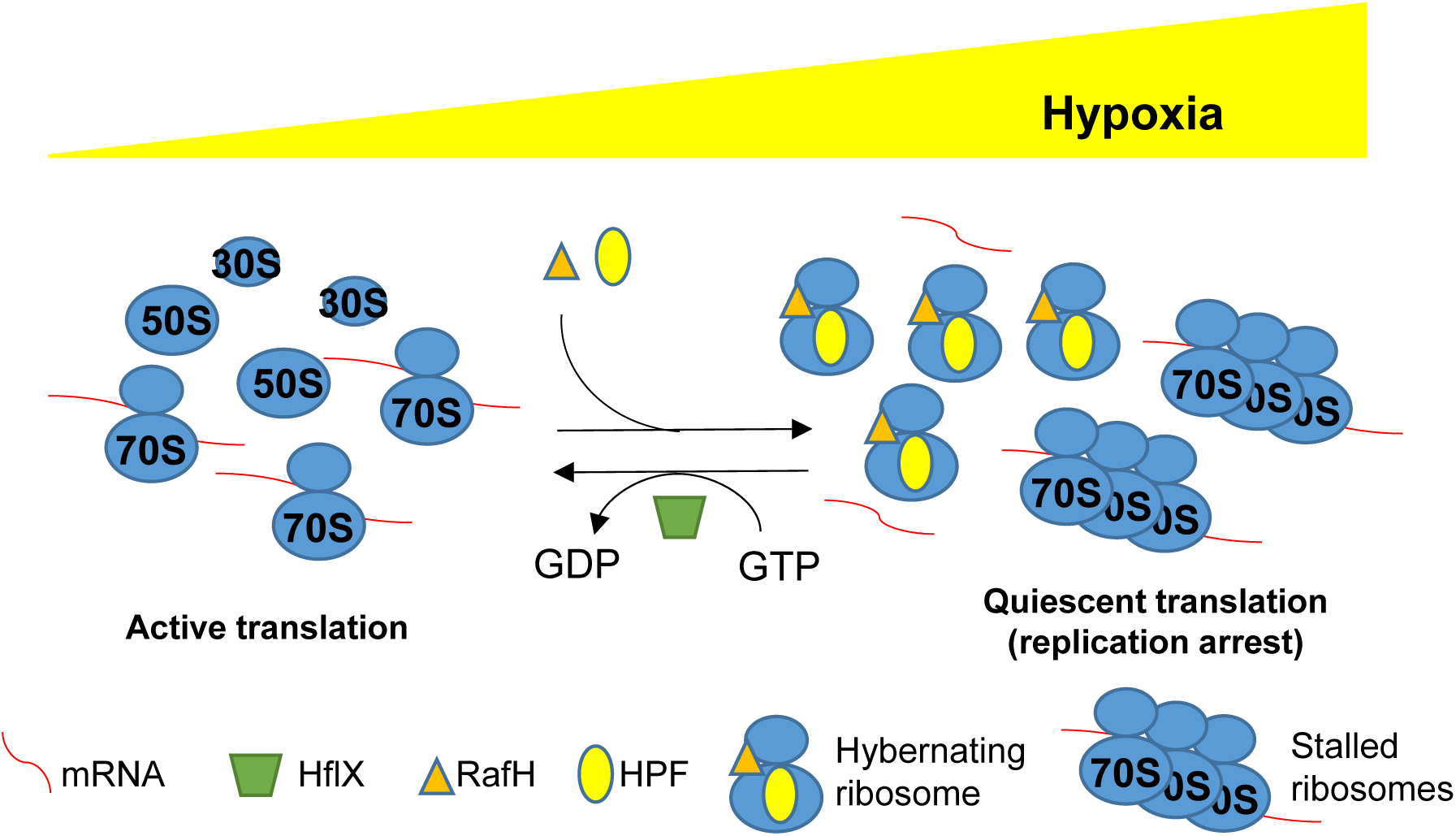
Illustration of the role of mycobacterial HflX in the hypoxic stress. As oxygen tension decreases, accumulation of stalled ribosomes and translation-incompetent hibernating ribosomes results in lower translational activity that eventually leads to bacterial replication arrest. By splitting stalled ribosomes and hibernating ribosomes, HflX controls the pool of translationally active ribosomes, thereby controlling the overall translational activity of the bacterium, and entry into the non-replicating state.

Overall, our work uncovers the physiological role of the highly conserved HflX GTPase in slow growing pathogenic mycobacteria, and provides further insights into the mechanisms by which this pathogen adapts to its environment. Such fundamental knowledge may help design alternative strategies to accelerate or potentiate the killing efficacy of current TB drugs.

## Methods

### Strains, Plasmids and Growth Conditions

Bacterial strains, plasmids, and primers used are listed in Appendix Tables S5 and S6. Mycobacteria were grown in Middlebrook 7H9 liquid medium (BD Difco™, CAT No.: 271310) supplemented with 0.5 % (v/v) glycerol, 0.05 % (v/v) Tween 80 and 10 % (v/v) Albumin and Dextrose; and plated on Middlebrook 7H11 Agar Base (BD Difco™, CAT No.: 271310) supplemented with Middlebrook OADC (BD Difco, CAT No.: 212351). When appropriate, hygromycin (50 μg/ml) (Roche, CAT No.: 10843555001) and kanamycin (50 μg/ml) (Thermo-Fisher Scentific, CAT No.: 11815032) was added to the media.

*E. coli* K-12 Δ*hflX* strain was purchased from Keio Collection (JW4131-1). *E. coli* strains were grown in Luria-Bertani (LB) broth (Sigma-Aldrich, CAT No.: L3022) and agar (BD Difco™, CAT No.: 244520). All pre-cultures were grown from frozen stock seeded in LB and cultured at 37 °C overnight under shaking conditions. Antibiotic ampicillin (100 μg/ml) (Sigma-Aldrich, CAT No.: A9518) was added into the medium where necessary for plasmid maintenance. Arabinose (Sigma-Aldrich, CAT No.: 10850) was added to the cultures when indicated for gene induction. Information on strains is provided in Appendix Table S5.

### Construction of knock-out and complemented strains

*M. bovis* BCG Δ*hflX* was generated via a double homologous recombination event, as previously described (Bardarov, 2002). Briefly, homologous regions (HR) flanking the 5’ and 3’ends of *hflX* were amplified from the WT BCG genome using primers described in Table S2, and cloned into the pYUB854 vector (Bardarov, 2002) with a hygromycin-resistance (*hygr*) cassette lies between both HRs. Transformants were plated onto 7H11 agar plates containing hygromycin, and resistant colonies were selected after 3 weeks of incubation for further validation. Successful knockouts were verified at the genomic level by PCR and at the transcriptional level by qRT-PCR.

The BCG Δ*hflX* mutant strain was complemented by reintroducing WT *hflX* open-reading frame (ORF) under the control of constitutive *hsp60* promoter (BCG Δ*hflX* :: *phflX*) back into the genome using pMV306 integrative plasmid (Stover, 1991). Transformants were plated onto kanamycin-7H11 agar plates, and resistant colonies were selected for further screening after 3 weeks incubation. Successfully complemented Δ*hflX* clones were validated at the genomic level by PCR and at the transcriptional level by qRT-PCR.

*E. coli* K-12 Δ*hflX* strain was complemented by reintroducing *hflX* ORF using pBAD replicative plasmid (Invitrogen, CAT No.: V44001) *E. coli hflX* locus was amplified from WT *E. coli* K-12 genome (*E. coli ΔhflX* :: *phflX*), while BCG *hflX* sequence was codon-optimized for *E. coli* expression (Stothard, 2000) and synthesized by GenScript (New Jersey, USA) (*E. coli ΔhflX* :: *pBCGhflX*). Both ORFs were expressed under the control of inducible arabinose promoter (*ara*), and the final plasmid vectors were electroporated into Δ*hflX E. coli*. Transformants were plated onto LB plates containing ampicillin. Resistant colonies were picked and verified at the transcriptional level by qRT-PCR.

### Heat Shock assay

Mid-log phase WT, Δ*hflX*, and *hflX* complemented *E. coli* cultures (OD_600nm_ 0.6) were diluted down to OD_600nm_ 0.1 with fresh LB broth. The bacterial cultures were then exposed to a temperature at 55°C for 10 minutes as previously described (Zhang et al., 2015). Bacterial viability was determined by plating on LB plates and incubated at 37°C, and the colony forming units (CFU) were enumerated. The same parameter of heat shock was applied to Δ*hflX*, and *hflX* complemented BCG cultures.

### Wayne model

The protocol was performed based on the previously described Wayne hypoxia model (Wayne, 1996). In short, mid-log phase BCG cultures (OD_600nm_ 0.6) were maintained in supplemented Dudos media and diluted to a final volume of 17mL (OD_600nm_ 0.005) in a glass tube containing a magnetic stir bar. The ratio between headspace and culture volume was also kept constant, as previously described (Wayne, 1996). The tubes were then tightly sealed with an airtight silicone seal cap and several layers of Parafilm M® to prevent oxygen diffusion. The tubes were incubated on a magnetic platform set at 170 rpm at 37°C for 3 weeks. Methylene blue was added at 0.015 µg/mL as a hypoxia control. At indicated time points (days 0, 3, 6, 8, 10, 14, 17 and 21), the airtight seal was broken open to measure turbidity (OD_600nm_) and enumerate CFU by plating appropriate dilutions of bacterial cultures on 7H11 agar plates to determine bacterial viability after hypoxic exposure.

To test antibitoics susceptibility, antibiotics [BDQ: Bedaquiline; INH: Isoniazid (Sigma-Aldrich, CAT No.: PHR1937); STM: Streptomycin (Sigma-Aldrich, CAT No.:S6501); RIF: Rifampicin (Sigma-Aldrich, CAT No.:R3501); CM: Chloramphenicol (MP-Bio, CAT No.:02190321); ETM: Ethambutol (Sigma-Aldrich, CAT No.:E4630)] were quickly injected into the tubes using a needle syringe, and the tubes were sealed back with several layers of parafilm. The bacterial cultures were incubated for another 5-days period on the magnetic platform with constant agitation at 100 rpm at 37 °C before plating onto 7H11 agar plates and incubation for 3 weeks at 37°C and 5% CO_2_. CFU were enumerated and compared to the drug-free control.

### Mice Infection

Animal experiments were approved by the Institutional Animal Care and Use Committee of the National University of Singapore (NUS) under protocol R16-0531 and were performed in the AALAAC-accredited animal facilities at NUS. Adult (7–8 weeks old) female Jackson C57BL/6 mice were purchased from InVivos (Singapore) and were intratracheally (IT) infected with ∼10^6^ CFU of *M. bovis* BCG strains (WT, Δ*hflX*, and Δ*hflX* :: *phflX*). At the indicated time points, lungs, lymph nodes, and spleens from euthanized mice were harvested and homogenized in PBS + 0.1% Triton X-100. Appropriate dilutions of the organ homogenates were plated onto 7H11 agar plates for CFU determination after 2 weeks incubation at 37°C and 5% CO_2_.

### Quantification of Intracellular ATP

A previously described method was followed (Rao, 2007). Briefly, intracellular ATP was quantified by using the BacTiter-Glo Microbial Cell Viability Assay Kit (Promega CAT No.: G8230). Aliquots of 100 µl of bacterial culture were collected at various time points and mixed with an equal volume of the BacTiter-Glo reagent and incubated for 5 min in the dark. The emitted luminescence was detected by using M200 Pro plate reader and was expressed as relative luminescence units.

### Measurement of the Membrane Potential

A previously described method was followed (Rao, 2007). Briefly, the membrane potential (ψ) was detected by using Baclight^TM^ Bacterial Membrane Potential Kit (Invitrogen™ CAT No.: B34950). 100 µl of bacterial cultures were collected at various time points and mixed with an equal volume of 60 µM DiOC2 (3,3-diethyloxa-carbocyanine iodine, fluorescent dye). After for 30 min at 37°C, the suspensions were analyzed using M200 Pro plate reader (Tecan Trading AG, Switzerland) at green fluorescence (ex: 488nm, em: 530nm) and red fluorescence (ex: 488nm, em: 630nm). Data were expressed as relative fluorescence units (RED/GREEN ratio). CCCP (Carbonyl cyanide *m*-chlorophenyl hydrazine) (Sigma-Aldrich, CAT No.:C2759) at a final concentration of 10 µM, was used as a positive control for membrane potential disruption.

### Turbidity-based growth inhibition assay

Mid-log *M. bovis BCG* cultures (OD_600nm_ 0.6) were diluted to OD_600nm_ 0.05 in 7H9 media. Bacterial suspensions were then dispensed in a transparent U-bottom 96-well plate (Greiner-Bio, CAT No.: 650180) (200 μL/well), containing 2-fold serially diluted antibiotic. The plates were incubated for 5 days at 37 °C. Bacterial suspensioms were manually resuspended before OD_600_ was measured using M200Pro plate reader (Tecan Trading AG, Switzerland). The minimum inhibition concentration 50 (MIC_50_), defined as the drug concentration that is required to inhibit 50% of bacterial growth (compared to drug-free control) was calculated.

### RNA extraction, cDNA synthesis, and qRT-PCR

Both *E. coli* and BCG cultures were treated with RNAprotect Cell Reagent (Qiagen, CAT No.:76526) before lysis. Treated bacterial cells were then centrifuged, and the pellet was resuspended in TE buffer with 20 mg/ml of lysozyme (Sigma-Aldrich, CAT No.:L7651) at room temperature for 20 minutes. BCG cultures had an additional disruption step using bead-beating before RNA extraction. The total RNA was isolated using the RNeasy®Mini Kit (Qiagen, CAT No.:74104), according to the manufacturer’s protocol. Extracted total RNA was further treated with TURBO DNA-*free*™ kit (Invitrogen™, CAT No.: AM1907) to remove genomic DNA contamination, according to the manufacturer’s protocol. RNA concentration and purity were determined using the NanoDrop 1000 Spectrophotometer™, and aliquots were stored at −80°C. Complementary DNA (cDNA) from extracted RNA was obtained using the iScript™ cDNA Synthesis Kit (Bio-rad, CAT No.:1708890) according to the manufacturer’s protocol. Relative mRNA abundance was then measured using the iTaq™ SYBR® Green Supermix (Bio-Rad, CAT No.:1725151) (refer to Table 2 for gene-specific primer sequences), and the 7500 Fast Real-Time PCR System (Applied Biosystems). Both the reaction mix and PCR cycling conditions followed the manufacturer’s instructions. The relative abundance of each *E. coli* and mycobacterial gene target was determined using house-keeping genes *rssA* or *sigA*, respectively, to normalize mRNA levels. The mRNA level of each target gene in the KO strain was expressed relative to the mRNA level measured in WT.

### Scanning electron microscopy (SEM)

Mycobacteria grown in the Wayne model were harvested at the indicated time points and coated on polylysine-coated glass coverslips, and fixed in 2.5% glutaraldehyde in 0.1 M phosphate buffer for 1 h (pH 7.4) at room temperature. The coverslips were then treated with 1% osmium tetroxide (Ted Pella Inc) at room temperature for 1 h, and then dehydrated through a graded ethanol series from 25% to 100% and critical point dried using a CPD 030 critical point dryer (Bal-Tec AG, Liechtenstein). The cell surfaces were coated with 15 nm of gold by sputter coating using a SCD005 high-vacuum sputter coater (Bal-Tec AG). The coated samples were examined with a field emission JSM-6701F Scanning Electron Microscope (JEOL Ltd., United States) at an acceleration voltage of 8 kV using the in-lens secondary electron detector.

### Tandem Mass Tag (TMT) mass spectrometry

Mycobacterial cultures was harvested at the indicated time points and washed twice with 1X PBS. Proteins were extracted using lysis buffer (8 M urea, 2 M thiourea, 4% CHAPS, 40mM DTT) supplemented with Halt protease inhibitor complete cocktail (Thermo Fisher) by bead beating (50 Hz for 3 five-minute cycles, TissueLyzer II (Qiagen) with 0.1 mm silica beads). Bead beating chambers were chilled at 4 °C before and between cycles. Extracts were centrifuged at 14,000 g for 15min at 4 °C and the supernatant was collected. Overnight trichloroacetic acid/acetone precipitation was performed with 2D Clean-Up kits (GE Healthcare) as instructed by the manufacturer. Air-dried protein pellets were resuspended in 10 mM triethylammonium bicarbonate (TEAB) buffer (pH 8.5) with 8M urea. Protein concentrations were determined by BCA assay (Thermo Science). Protein quality and quantities were checked by SDS-PAGE electrophoresis (12% polyacrylamide gels) and UV spectrometry (Nanodrop, Thermo Scientific). A total of 100 μg protein from each condition was subjected to in-solution trypsin digestion before labelling the resultant tryptic peptides using the TMT-6plex Isobaric Label Reagent Set (Thermo Scientific, Rockford, IL, USA) according to the manufacturer’s protocol. The labeled samples were combined prior to fractionation using a high pH reverse phase HPLC on a Xbridge™ C18 column (4.6 × 250 mm, Waters, Milford, MA, USA) and subsequent analysis by LC-MS/MS.

The fractionated peptides were separated and analyzed using a Dionex Ultimate 3000 RSLCnano system coupled to Q Exactive tandem mass spectrometry (Thermo Fisher Scientific, MA, USA). Separation was performed on a Dionex EASY-Spray 75 μm × 10 cm column packed with PepMap C18 3 μm, 100 Å (Thermo Fisher Scientific) using solvent A (0.1% formic acid) and solvent B (0.1% formic acid in 100% ACN) at flow rate of 300 nL/min with a 60 min gradient. Peptides were then analyzed on the Q Exactive apparatus with the EASY nanospray source (Thermo Fisher Scientific) at an electrospray potential of 1.5 kV. A full MS scan (350–1,600 m/z range) was acquired at a resolution of 70,000 and a maximum ion accumulation time of 100 ms. Dynamic exclusion was set as 30 s. The resolution of the higher energy collisional dissociation (HCD) spectra was set to 350,00. The automatic gain control (AGC) settings of the full MS scan and the MS2 scan were 5E6 and 2E5, respectively. The 10 most intense ions above the 2,000 count threshold were selected for fragmentation in HCD, with a maximum ion accumulation time of 120 ms. An isolation width of 2 m/z was used for MS2. Single and unassigned charged ions were excluded from MS/MS. For HCD, the normalized collision energy was set to 30. The underfill ratio was defined as 0.3%. Raw data files from the three technical replicates were processed and searched using Proteome Discoverer 2.1 (Thermo Fisher Scientific). The raw LC-MS/MS data files were loaded into Spectrum Files (default parameters set in Spectrum Selector) and TMT 6-plex was selected for the Reporter Ion Quantifier. The SEQUEST HT algorithm was then used for data searching to identify proteins using the following parameters; missed cleavage of two; dynamic modifications were oxidation (+15.995 Da) (M) and deamidation (+0.984 Da) (NQ). The static modifications were TMT-6plex (+229.163 Da) (any N-terminus and K) and Carbamidomethyl (+57.021 Da) (C). The false discovery rate for protein identification was <1%. The Normalization mode was set based on total peptide amount.

### Cloning, expression and purification of recombinant HflX proteins

*M. tuberculosis* (Mtb) *hflX* gene sequence encoding amino acids 1-435 was cloned into a pNIC-CH2 expression vector with a His_6_ tag at its C-terminus by Protein Production Platform (PPP), NTU. pNIC-CH2 HflX was mutated with PCR-based mutagenesis to produce HflX AAY expression plasmid. *E. coli* HflX expression plasmid was also generated by PPP, NTU. Mtb HflX, Mtb HflX AAY, and *E.Coli* HflX constructs were transformed into BL21(DE3)-T1^R^ competent cells (Sigma-Aldrich, CAT No.:B2935) for protein expression. For protein expression, 10 mL of overnight starter bacterial culture was added to 1L of LB media supplemented with kanamycin and chloramphenicol and cultured at 37°C on a shaker to OD_600_ of 0.8 prior to addition of 0.5 mM IPTG and overnight incubation at 18°C. After centrifugation, the bacterial pellet was resuspended in cold lysis buffer (100 mM Na Hepes, pH 7.5, 500 mM NaCl, 10 mM imidazole, 1 mM TCEP, and 10% glycerol), and lysed using LM20 microfluidizer with a pressure of 20,000 psi. Clarified lysates were collected after centrifugation for three-step purification, including nickel affinity chromatography, ion-exchange chromatography, and size exclusion chromatography. The protein was eluted in gel filtration buffer (20 mM Hepes, pH 7.5, 300 mM NaCl, 1 mM TCEP,10% glycerol) and concentrated using Vivaspin turbo with a 10 kDa molecular mass cutoff concentrator (Sartorius) to a final concentration of 1 mg/mL. Protein quality and purity were assessed by SDS-PAGE, and the suspensions were stored at −80°C.

### Generation of anti-HflX monoclonal antibody

BALB/c mice (females, 6 weeks old) were injected intraperitoneally with 25 µg of purified Mtb HflX protein as described above, mixed with incomplete Freund’s adjuvant (Sigma-Aldrich, USA) in a 1:1 volumetric ratio for three cycles at 2-week intervals. A final booster immunization consisted of administering intravenously 25 µg of the same antigen without adjuvant. Three days later, the splenocytes from the euthanized BALB/c mice were obtained and fused with myeloma cells NS-1 using standard hybridoma methods (Köhler, 1975; Yokoyama, 2013). Screening of hybridoma cells and titer analysis were carried out as described previously (Köhler, 1975; Yokoyama, 2013). The monoclonal antibody selected was confirmed to detect Mtb/BCG HflX in an ELISA assay and by dot blot, but was unable to detect HflX in Western blot, indicating that this antibody likely recognized a conformational epitope.

### Immunoprecipitation and LC/MS

Mid-log BCG WT and KO cultures (7H9) were harvested and the bacteria pellets were washed twice with 1X PBS, before proceeding to protein extraction. Bacterial lysates were suspended in Pierce^TM^ IP lysis buffer (Thermo Fisher Scientific, CAT No.:87787) and supplemented with 1X EDTA and 1X Halt Protease inhibitor cocktail (Thermo-Scientific, CAT No.:78440) and were lysed using bead beating (50 Hz for 3 five-minute cycles, TissueLyzer II (Qiagen) with 0.1 mm silica beads). 800 μg of lysates were then co-incubated with monoclonal anti-HflX that was pre-treated with Dynabeads Protein G (Thermo Fisher Scientific, CAT No.:1004D) for immunoprecipitation. The dynabeads were washed 3X with 1X PBS and 0.05% Tween20. Co-IP elutes were extracted with 1X NUPAGE^TM^ LDS sample buffer (Thermo Fisher Scientific, CAT No.: N0007). The proteins were separated on an 8–20% gradient SDS-PAGE and subjected to in-gel digestion. The peptides were separated and analyzed using a Dionex Ultimate 3000 RSLCnano system coupled to a Q Exactive instrument as described in the Tandem Mass Tag (TMT) mass spectrometry section above. Raw data files were converted to mascot generic file format using Proteome Discoverer 1.4 (Thermo Fisher Scientific). The Mascot algorithm was then used for data searching to identify proteins. emPAI value reported by Mascot was used for label free protein quantitation and proteins identified in the WT only (after minus the background proteins identified in the hflX KO strain) were shortlisted for further analysis.

### GTPase/ATPase hydrolysis assay

GTP/ATPase hydrolysis was quantified using malachite green phosphate assay kit (Sigma-Aldrich, CAT No.: MAK307) according to the manufacturer protocol. Briefly, a 300 µL of reaction mixture was prepared consisting of the respective purified proteins at a final concentration of 1 µM, 300 µM GTP or ATP and reaction buffer (50 mM Tris-HCl, pH 8.0, 200 mM NaCl, 1 mM DTT and 5 mM MaCl_2_). 80 µL of the reaction mixture were collected at 1 hr, 3 hr, and overnight respectively, and mixed with 20 µL of malachite green phosphate assay reagent. After 5 minutes incubation at room temperature for 5 min, OD_620nm_ was measured with a Tecan multimode microplate reader (Tecan Trading AG, Switzerland).

### Isothermal titration calorimetry (ITC) assay

ITC assay was performed to evaluate the direct interaction between HflX protein with ligands involved in GTPase hydrolysis, including GDP, GTP, and GMP-PNP. Both HflX protein and the ligands were prepared in (20 mM Hepes, pH 7.5 and 300 mM NaCl). Briefly, 800 µM of GDP, GTP or GMP-PNP was loaded into the syringe while 100 µM of Mtb HflX protein was loaded into the experimental cell. Titrations were performed at 25 °C consisting of an initial injection at 0.5 µL and 19 injections at 2 µL of GDP, GTP or GMP-PNP into Mtb HflX protein until saturation was reached. Thermodynamic data were analyzed with a single-site fitting model using MicroCal PEAQ-ITC analysis software provided by the manufacturer.

### Ribo-sequencing

*Extraction of RNA and ribosomes-* Mycobacterial cultures grown in the Wayne model were harvested at day 8 and treated with 100 µg/mL Chloramphenicol (Sigma Aldrich, CAT No.: C0378) for 3 min before centrifugation at 4,500 g for 10min at 4 °C followed by one wash with chilled 1X polysome buffer (20mM Tris HCL, 100mM NaCl, 5mM MgCl_2_, 100 µg/mL Chloramphenicol). Cell pellets were resuspended in ice-cold lysis buffer (1X polysome buffer, 1% Triton X 100, 1mM DTT, 20 U/mL Turbo DNAse, 0.1% NP40, 100 µg/mL Chloramphenicol) and flash frozen in liquid nitrogen. The frozen cells were pulverized using Cell Crusher tissue pulverizer (Cell Crusher, CAT No.: 607KSL) according to the manufacturer’s protocol. The grinding jar was pre-chilled in liquid nitrogen. The extracts were centrifuged at 14,000 g for 20min at 4 °C and the supernatant was collected. RNA concentration was determined by Qubit^TM^ RNA HS assay kit (Thermo Fisher, CAT No.: Q32852).

*Ribosome profiling-* Pulverized cells were thawed and the soluble cytoplasmic fraction was isolated by centrifugation at top speed for 20 min at 4 °C (Oh, 2011). Supernatant was collected and the clarified lysates were digested with RNase I for 1h at room temperature. Digestion was stopped with SuperaseIN and monosomes purified by size exclusion chromatography on MicroSpin S-400 HR columns (GE Healthcare) as described (Shamimuzzaman, 2018). Size selection of footprints with length 15-40 nt was performed by electrophoresis on 15% TBE-urea gels. 3’ termini of ribosome footprints were dephosphorylated with T4 polynucleotide kinase. Illumina ready RIBO-seq libraries were prepared using SMARTer smRNA kit (TakaraBio). Library concentrations were measured by Qubit fluorometer and their quality assessed on a Agilent 2100 bioanalyzer. RIBO-seq libraries were sequenced on a Illumina Novaseq 6000 sequencer.

*RNA-sequencing-* To obtain matched RNA-seq libraries, total RNA was purified from an aliquot of cell lysate, rRNA was depleted using RIBO-Minus transcriptome isolation kit (Invitrogen) following manufacturer’s instructions. mRNA fragmentation was conducted for 25 min at 94 °C to generate RNA fragments of similar sizes as those of the ribosome footprints. SMARTer smRNA kit (TakaraBio) was used to generate Illumina ready RNA-seq libraries. Library concentrations were measured by Qubit fluorometer and their quality assessed on a Agilent 2100 bioanalyzer. RNA-seq libraries were sequenced on a Illumina Novaseq 6000 sequencer.

### Loebel nutrient starvation model

The starvation model follows a previously described method (Betts, 2002; Loebel, 1933). Briefly, mid-log phase (OD_600nm_ 0.6) *M. bovis* BCG cultures grown in 7H9 were washed thrice with sterile DPBST (1 X PBS supplemented with 100 mg/L CaCl_2_ and 100 mg/L MgCL_2_-6H_2_O and 0.05% Tween 80). The pellet was then resuspended in 50 mL sterile at an OD_600nm_ of 0.1 within a one-liter roller bottle (Corning® Roller Bottles, Tissue Culture Treated, 490 cm2, cap plug seal, CAT No..: CLS430195-40EA). The cultures were then incubated at 37 °C on a rolling platform for 2 weeks. At the indicated time points (days 0, 6, 9, 14), turbidity was measured at OD_600nm_, and CFU were enumerated onto 7H11 agar plates.

### THP-1 macrophage infection assay

THP-1 cells (American Type Culture Collection) were grown in RPMI 1640 (Gibco CAT No.: 22400-15) supplemented with 10% fetal bovine serum (Gibco CAT No.: 10270-106), 0.01 mM sodium pyruvate (Gibco CAT No.: 11360-070), 1% Glutamax (Gibco CAT No.: 35050-061) and 0.5 µM Beta-mecaptoethanol (Gibco CAT No.: 21985-023). THP-1 were seeded in 24-wells plate at 5 X 10^4^ cells/well. THP-1 cells differentiated with 100 ng/mL phorbol-12-myristate 13-acetate (PMA) (Sigma-Aldrich, CAT No.:79346) were allowed to adhere for 24 hours before infection. Dispersed bacilli were incubated with differentiated THP-1 cells at a multiplicity of infection (MOI) of 2 for 1h at 37°C, 5% CO_2_. The cells were then washed thoroughly twice with pre-warmed PBS and then incubated for 1 to 5 days at 37 °C and 5% CO_2_. After incubation, cells were lysed, and bacteria were harvested and plated on 7H11 agar plate for CFU enumeration 3 weeks later.

### Computational modeling

Web-based protein modeling platform Phyre 2.0 was used (Kelley, 2015). The Mtb H37Rv HflX primary protein sequence was obtained from the NCBI Gene database and entered into the Phyre 2.0 with the modeling mode set to “Intensive”. Homology modeling uses the *E. coli* HflX crystal structure (PDB entry: 5ADY) as a template to produce the model. Model homology was individually assessed based upon the parameters set for each platform. Phyre 2.0 utilizes a Confidence score based on HHsearch, which uses a profile hidden Markov model to assess the quality of model alignment with the template (Soding, 2005).

### Statistical analysis

Statistical analyses were generated from Prism 7.0 (GraphPad, USA) and tests used are indicated in the figure legends. One-way and two-way ANOVA were conducted on experiments comparing across different groups under single and multiple conditions, respectively, with Bonferroni correction as *post-hoc* test. Results with *p*-values <0.05 were defined as statistically significant.

## Acknowledgements

This work was supported by a grant from the Ministry of Education (Singapore) allocated to SA. We would like to thank the Antibody Core Facility at the Life Sciences Institute, for their assistance in generating the anti-HflX monoclonal antibody; and TB-SEQ, Inc (Palo Alto, USA) for performing library preparation, ribosome sequencing and bioinformatics services. We would also like to thank Dr. Rohan Williams from SCELSE (NUS) for his insightful comments on data analyzing.

## Author contributions

-NGAN Jie Yin Grace, PAUNOOTI Swathi, PETHE Kevin, SZE Siu Kwan, LESCAR Julien, and ALONSO Sylvie designed the experiments, and analyzed the data.

-NGAN Jie Yin Grace, PAUNOOTI Swathi, MENG Wei, NG Sze Wai, JAAFA Muhammad Taufiq, JIA Huan, CHO Su Lei Sharol, LIM Jieling, KOH Hui Qi Vanessa, ABDUL GHANI Noradibah, performed the experiments.

-TSE Wilford, NGAN So Fong Cam, analyzed data.

-NGAN Jie Yin Grace, ALONSO Sylvie wrote the manuscript.

## Appendix Figure Legends

**Figure S1.**
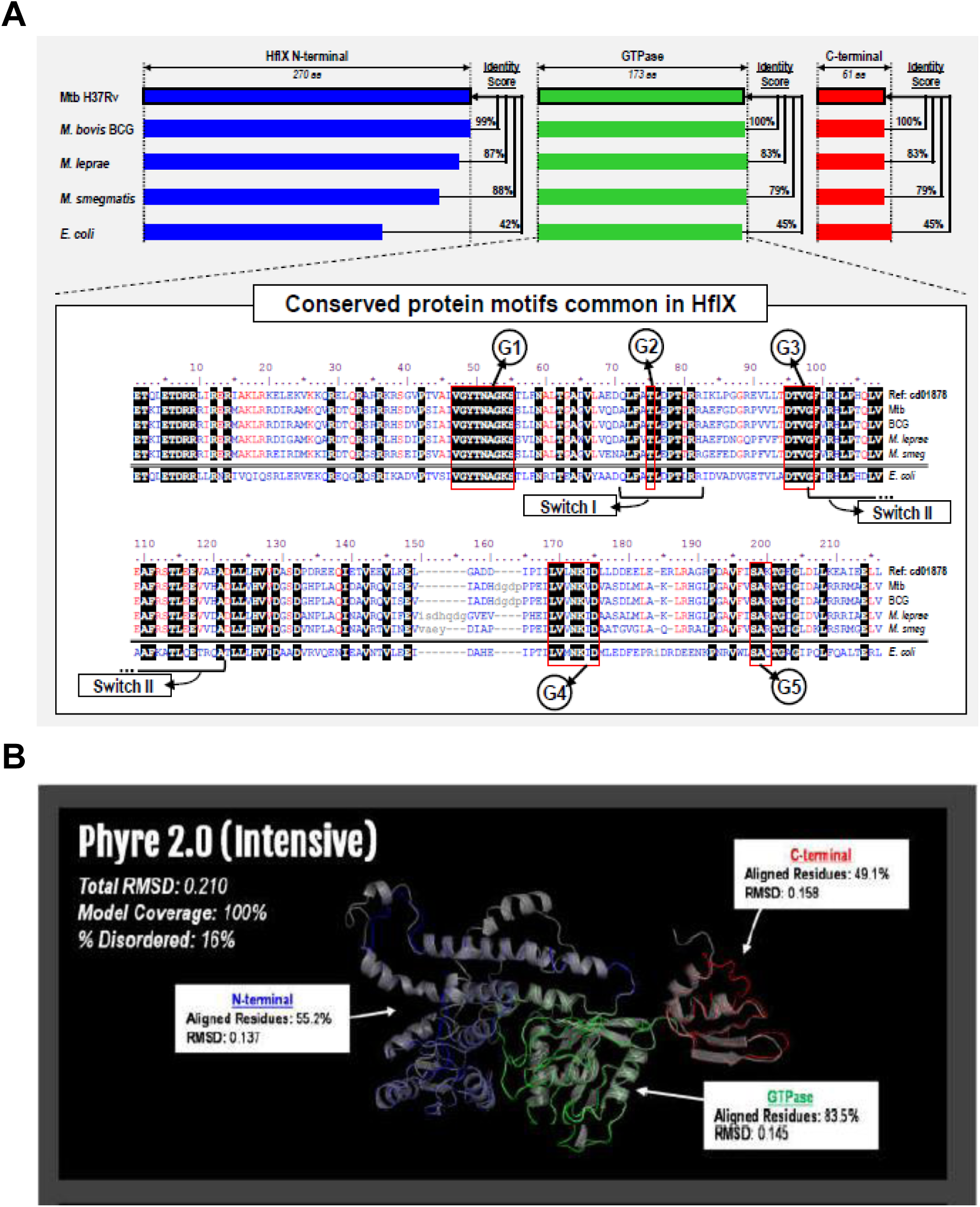
A. Protein sequence alignment of mycobacterial and *E. coli* HflX. Protein motifs important for GTPase function were also identified (red boxes). G1–5 boxes (circled), Switch I and II regions (boxed). B. Computational modelling of Mtb H37Rv HflX superimposed with *E. coli* HflX crystal structure. Model quality was determined by the percentage of residues that were successfully modelled (Model Coverage). Alignment quality for complete models and their individual domains were quantified by an RMSD score where 0 indicates perfect alignment.

**Figure S2.**
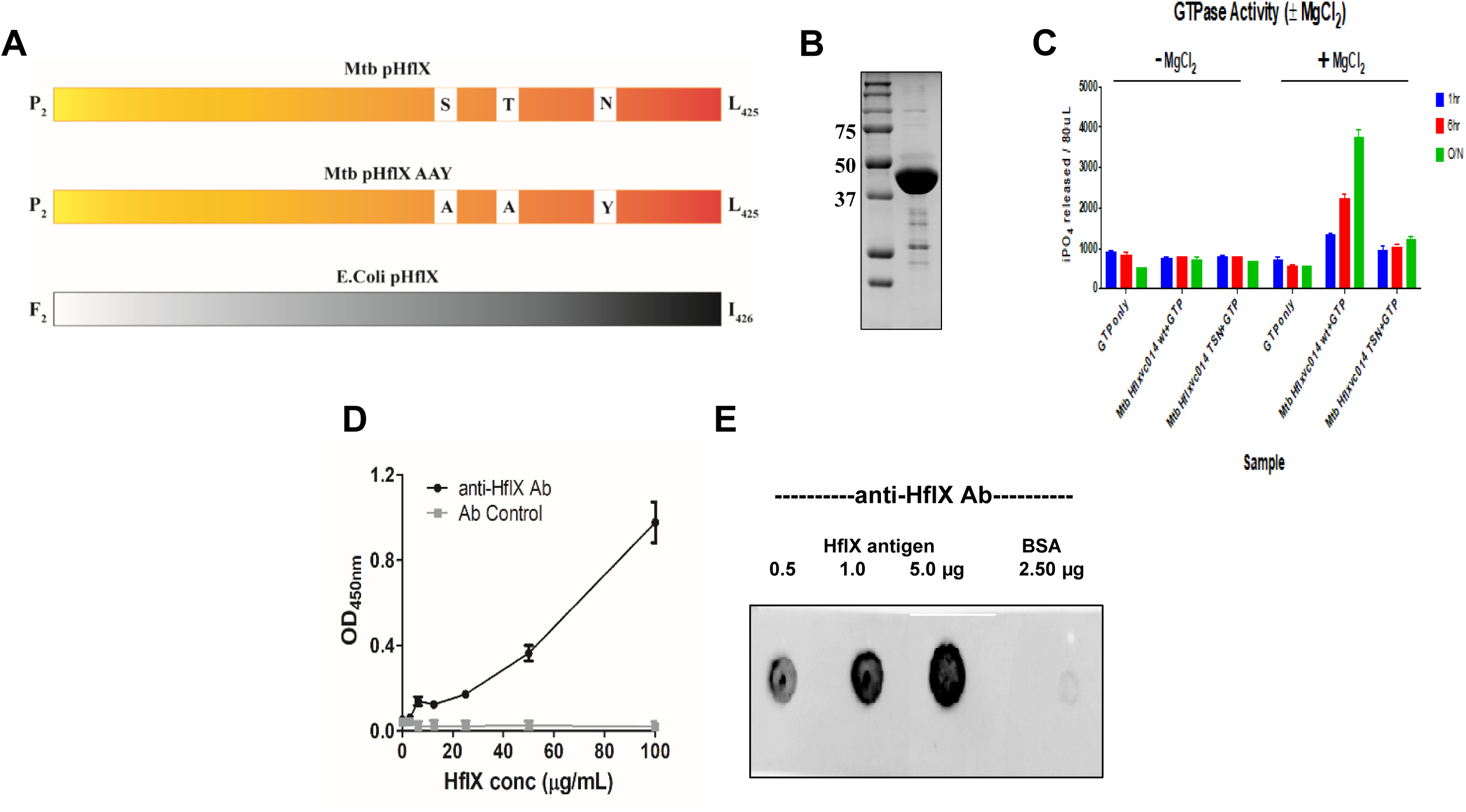
A. Schematic illustration of Mtb pHflX, abrogated GTPase Mtb pHflX AAY and E*. coli* HflX. B. Coomassie gel showing purified Mtb pHflX. C. GTPase activity of purified Mtb HflX in the presence or absence of magnesium ion. D. ELISA validation of the anti-HflX monoclonal Ab with coated HflX antigen. Data show mean ± SD of two independent experiments. E. Dot blot validation of anti-HflX monoclonal Ab detecting purified Mtb HflX.

**Figure S3.**
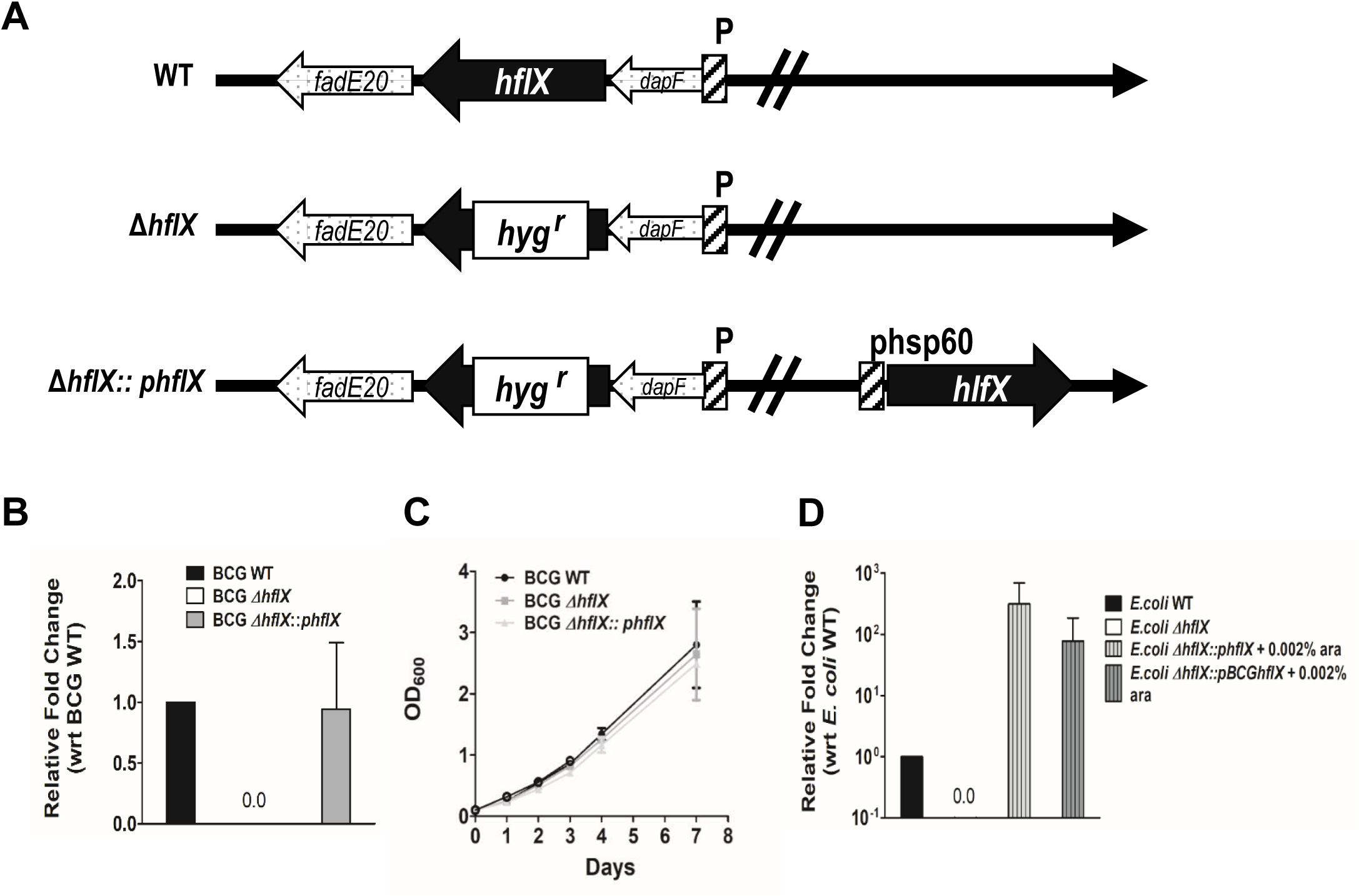
A. Schematic representation of the genetic construct of BCG Δ*hflX* and complemented strains as described in the methods. B. Expression level of *hflX* in BCG WT, Δ*hflX* and complemented strains determined by RT-PCR. Data show mean ± SD of two independent experiments. C. Growth profile of BCG WT, Δ*hflX* and complemented strains grown in 7H9 medium (normoxia). Data show mean ± SD of two independent experiments. D. Expression level of *hflX* determined by RT-PCR in *E. coli* WT, Δ*hflX* and Δ*hflX* complemented with homologous HflX or codon-optimized BCG HflX.

**Figure S4.**
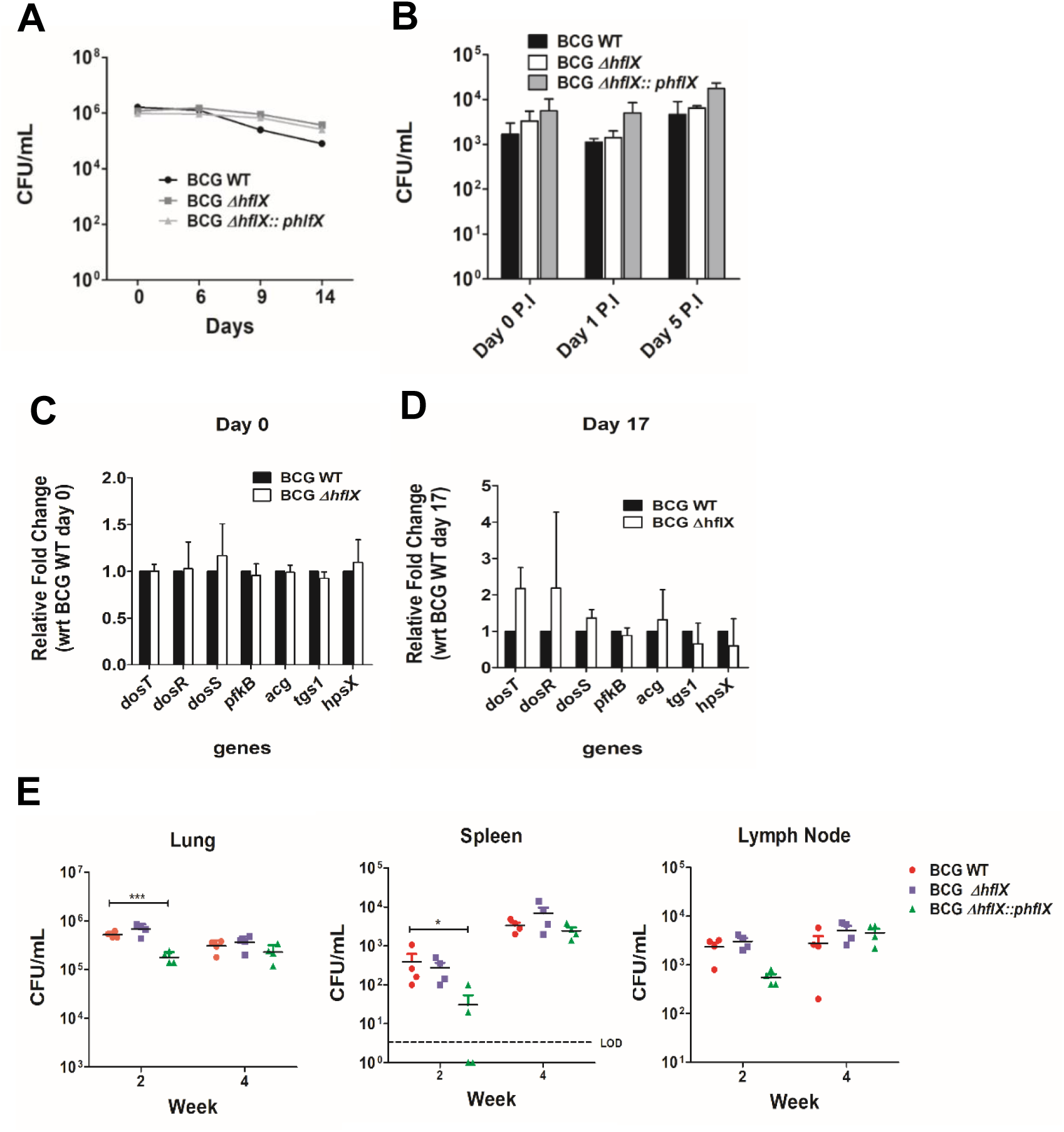
A. CFU from BCG WT, Δ*hflX* and complemented strains grown in Loebel starvation model as described in the methods. Data show mean ± SD of two independent experiments. B. CFU from THP-1 macrophage infected with BCG WT, Δ*hflX* and complemented strains (MOI: 2) as described in the methods. Data show mean ± SD of three independent experiments. C-D. Expression of *dos* regulon genes determined by RT-PCR in BCG WT and Δ*hflX* on day 0 (C) or day 17 (D) of the Wayne model. Data show mean ± SD of two independent experiments. E. CFU counts from C57BL/6 mice infected with BCG WT, Δ*hflX* and complemented strains. Organs were harvested at week 2 and 4 post-infection. LOD, limit of detection. Data information: All data show mean ± SD, *P < 0.05. Panels (E) one-way ANOVA with Bonferroni post-test.

**Figure S5.**
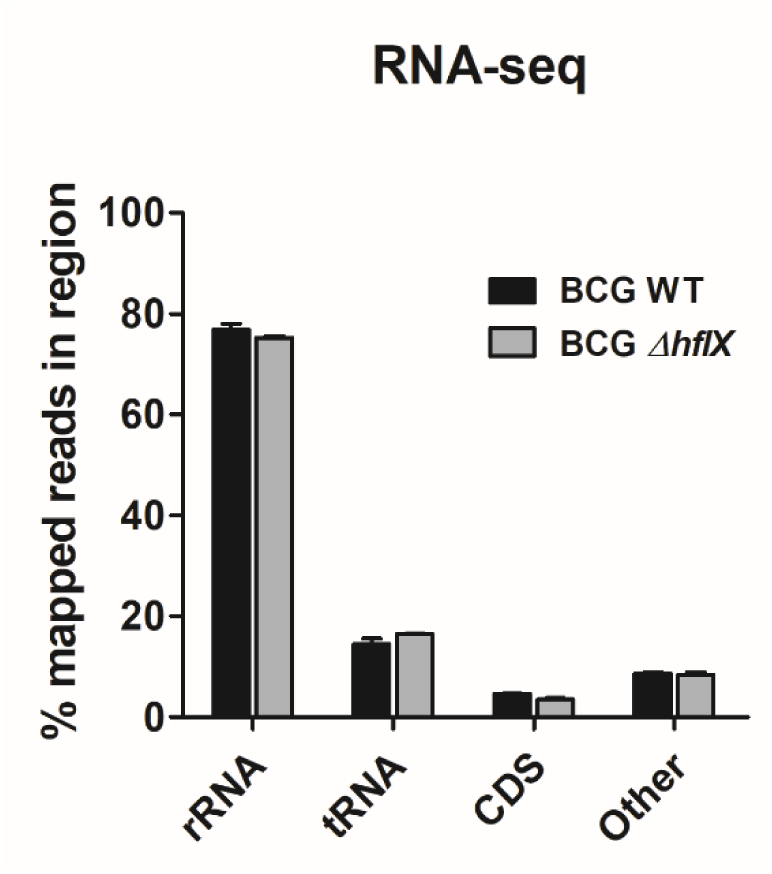
RNA sequencing data showing the percentage of mapped reads to respective regions of BCG Δ*hflX* compared to WT on day 8 (NRP-1) of Wayne model. CDS: coding sequence; Other: 5’ and 3’ untranslated regions (UTR). Data show mean ± SD of two independent experiments.

**Appendix Table S1.** Minimum Inhibitory Concentration of drugs against BCG WT, ΔHflX and complemented strains grown in 7H9 medium (normoxia).

| Strains | MIC50 (μM) |  |  |  |  |  |  |
| --- | --- | --- | --- | --- | --- | --- | --- |
|  | BDQ | INH | RIF | ETM | STM | CM | ERT |
| **BCG WT** | 0.05 | 1.25 | 0.1 | 8 | 1 | 25 | 3.125 |
| **BCG Δ*hflX*** | 0.06 | 1.25 | 0.1 | 6.25 | 1.25 | 30 | 6.25 |
| **BCG Δ*hflX* :: *phflX*** | 0.05 | 1.25 | 0.06 | 6.25 | 1 | 25 | 4 |

BDQ: Bedaquiline; INH: Isoniazid; RIF: Rifampicin; ETM: Ethambutol; STM: Streptomycin; CM: Chloramphenicol; ERT: Erythromycin

## References

Andini N, Nash, KA. (2006) Intrinsic macrolide resistance of the Mycobacterium tuberculosis complex is inducible. Antimicrob Agents Chemother 50: 2560–2562

Andreu N, Gibert, I. (2008) Cell population heterogeneity in Mycobacterium tuberculosis H37Rv. Tuberculosis (Edinb) 88: 553–559

Andries K, Verhasselt, P., Guillemont, J., Göhlmann, HWH., Neefs, JM., Winkler, H., Gestel, JV., Timmerman, P., Zhu, M., Lee, E., Williams, P., de Chaffoy, D., Huitric, E., Hoffner, S., Cambau, E., Truffot-Pernot, C., Lounis, N., Jarlier, V. (2005) A diarylquinoline drug active on the ATP synthase of Mycobacterium tuberculosis. Science 307: 223–227

Azad A, Sirakova, TD., Fernandes, ND., Kolattukudy, PE. (1997) Gene knockout reveals a novel gene cluster for the synthesis of a class of cell wall lipids unique to pathogenic mycobacteria. J Biol Chem 272: 16741–16745

Bagchi G, Chauhan, S., Sharma, D., Tyagi, JS. (2005) Transcription and autoregulation of the Rv3134c-devR-devS operon of Mycobacterium tuberculosis Microbiol 151: 4045–4053

Bardarov S, Bardarov, S., Pavelka, MS., Sambandamurthy, V., Larsen, M., Tufariello, JA., Chan, J., Hatfull, G., Jacobs, WR. (2002) Specialized transduction: an efficient method for generating marked and unmarked targeted gene disruptions in Mycobacterium tuberculosis, M. bovis BCG and M. smegmatis. Microbiology 148: 3007–3017

Basu A, Yap, MNF. (2017) Disassembly of the Staphylococcus aureus hibernating 100S ribosome by an evolutionarily conserved GTPase. PNAS 114: 8165–8173

Bennison D, Irving, SE., Corrigan, RM. (2019) The Impact of the Stringent Response on TRAFAC GTPases and Prokaryotic Ribosome Assembly. Cells 8: 1313

Betts J, Lukey, PT., Robb, LC., McAdam, RA., Duncan, K. (2002) Evaluation of a nutrient starvation model of Mycobacterium tuberculosis persistence by gene and protein expression profiling. Mol Microbiol 43: 717–731

Binder A, Adjemian, J., Olivier, KN., Prevots, DR. (2013) Epidemiology of nontuberculous mycobacterial infections and associated chronic macrolide use among persons with cystic fibrosis. Am J Respir Crit Care Med 188: 807–812

Bloom B, Murray, CJ. (1992) Tuberculosis: commentary on a reemergent killer. Science 257: 1055–1064

Boon C, Dick, T. (2002) Mycobacterium bovis BCG response regulator essential for hypoxic dormancy. J Bacteriol 184: 6760–6767

Boshoff H, Myers, TG., Copp, BR., McNeil, MR., Wilson, MA., Barry, CE 3rd. (2004) The transcriptional responses of Mycobacterium tuberculosis to inhibitors of metabolism: novel insights into drug mechanisms of action. J Biol Chem 279: 40174–40184

Bourne H, Sanders, DA., McCormick, F. (1990) The GTPase superfamily: a conserved switch for diverse cell functions. Nature 348: 125–132

Bretl D, Demetriadou, C., Zahrt, TC. (2011) Adaptation to Environmental Stimuli within the Host: Two-Component Signal Transduction Systems of Mycobacterium tuberculosis. Microbiol Mol Biol Rev 75: 566–582

Britton R (2009) Role of GTPases in bacterial ribosome assembly. Annu Rev Microbiol 63: 155–176

Britton R, Chen, SM., Wallis, D., Koeuth, T., Powell, BS., Shaffer, LG., Largaespada, D., Jenkins, NA., Copeland, NG., Court, DL., Lupski, JR. (2000) Isolation and preliminary characterization of the human and mouse homologues of the bacterial cell cycle gene era. Genomics 67: 78–82

Britton R, Powell, BS., Dasgupta, S., Sun, Q., Margolin, W., Lupski, JR., Court, DL. (1998) Cell cycle arrest in Era GTPase mutants: a potential growth rate-regulated checkpoint in Escherichia coli. Mol Microbiol 27: 739–750

Burian J, Ramón-García, S., Sweet, J., Gómez-Velasco, A., Av-Gay, Y., Thompso, CJ. (2012) The Mycobacterial Transcriptional Regulator whiB7 Gene Links Redox Homeostasis and Intrinsic Antibiotic Resistance. J Bio Chem 287: 299–310

Buriánková K, Doucet-Populaire, F., Dorson, O., Gondran, A., Ghnassia, JC., Weiser, J., Pernodet, JL. (2004) Molecular basis of intrinsic macrolide resistance in the Mycobacterium tuberculosis complex. Antimicrob Agents Chemother 48: 143–150

Caldon C, March, PE. (2003) Function of the universally conserved bacterial GTPases. Curr Opin Microbiol 6: 135–139

Campbell T, Brown, ED. (2008) Genetic interaction screens with ordered overexpression and deletion clone sets implicate the Escherichia coli GTPase YjeQ in late ribosome biogenesis. J Bacteriol 190: 2537–2545

Carreau A, El Hafny-Rahbi, B., Matejuk, A., Grillon, C., Kieda, C. (2011) Why is the partial oxygen pressure of human tissues a crucial parameter? Small molecules and hypoxia. . J Cell Mol Med 15: 1239–1253

Cladière L, Hamze, K., Madec, E., Levdikov, VM., Wilkinson, AJ., Holland, IB., Séror, SJ. (2006) The GTPase, CpgA(YloQ), a putative translation factor, is implicated in morphogenesis in Bacillus subtilis. Mol Genet Genomics 275: 409–420

Coatham ML, Brandon, H. E., Fischer, J. J., Schümmer, T. & Wieden, H.-J. (2016) The conserved GTPase HflX is a ribosome splitting factor that binds to the E-site of the bacterial ribosome. Nucleic Acids Research 44: 1952–1961

Dannenberg AJ (1993) Immunopathogenesis of pulmonary tuberculosis. Hosp Pract (Off Ed) 28: 51–58

Davies J, Davies, D. (2010) Origins and evolution of antibiotic resistance. Microbiol Mol Biol Rev 74: 417–433

Dey S, Biswas, C., Sengupta, J. (2018) The universally conserved GTPase HflX is an RNA helicase that restores heat-damaged Escherichia coli ribosomes. J Cell Bio 217: 2519–2529

Dheda KB, H., Huggett, JF., Johnson, MA., Zumla, A., Rook, GA. (2005) Lung remodeling in pulmonary tuberculosis. J Infect Dis 192: 1201–1205

Dutta D, Bandyopadhyay, K., Datta, A. B., Sardesai, A. A. & Parrack, P. (2009) Properties of HflX, an enigmatic protein from Escherichia coli. J Bacteriol 191: 2307–2314

Dutta N, Mehra, S., Didier, PJ., Roy, CJ., Doyle, LA., Alvarez, X., Ratterree, M., Be, NA., Lamichhane, G., Jain, Sk., Lacey, MR., Lackner, AA., Kaushal, D. (2010) Genetic requirements for the survival of tubercle bacilli in primates. J Infect Dis 201: 1743–1752

Dye C, Scheele, S., Dolin, P., Pathania, V., Raviglione, MC. (1999) Consensus statement. Global burden of tuberculosis: estimated incidence, prevalence, and mortality by country. WHO Global Surveillance and Monitoring Project. JAMA 282: 677–686

El-Sharoud W (2004) Ribosome inactivation for preservation: concepts and reservations. Sci Prog 87: 137–152

Fischer J, Coatham, ML., Bear, SE., Brandon, HE., De Laurentiis, EI., Shields, MJ., Wieden, H. (2012) The ribosome modulates the structural dynamics of the conserved GTPase HflX and triggers tight nucleotide binding. J Biochimie 94: 1647–1659

Foti J, Persky, NS., Ferullo, DJ., Lovett, ST. (2007) Chromosome segregation control by Escherichia coli ObgE GTPase. Mol Microbiol 65: 569–581

Franzblau S, DeGroote, MA., Sang, HC., Andries, K., Nuermberger, E., Orme, IM., Mdluli, K., Angulo-Barturen, I., Dick, T., Dartois, V., Lenaerts, AJ. (2012) Comprehensive analysis of methods used for the evaluation of compounds against Mycobacterium tuberculosis. Tuberculosis 92: 453–488

Fu L, Tai, SC. (2009) The Differential Gene Expression Pattern of Mycobacterium tuberculosis in Response to Capreomycin and PA-824 versus First-Line TB Drugs Reveals Stress- and PE/PPE-Related Drug Targets. Int J Micro 2009: 1–9

Gautam U, Sikri, K., Vashist, A., Singh, V., Tyagi, JS. (2014) Essentiality of DevR/DosR interaction with SigA for the dormancy survival program in Mycobacterium tuberculosis. J Bacteriol 196: 790–799

Gohara D, Yap, MF. (2018) Survival of the drowsiest: the hibernating 100S ribosome in bacterial stress management. Curr Genet 64: 753–760

Gold B, Nathan, C. (2017) Targeting Phenotypically Tolerant Mycobacterium tuberculosis. Microbiol Spectr 5: 10.1128

Gollop N, March, PE. (1991) A GTP-binding protein (Era) has an essential role in growth rate and cell cycle control in Escherichia coli. J Bacteriol 173: 2265–2270

Gomez J, McKinney, JD. (2004) M. tuberculosis persistence, latency, and drug tolerance. Tuberculosis (Edinb) 84: 29–44

Haagsma A, Podasca, I., Koul, A., Andries, K., Guillemont, J., Lill, H., Bald, D. (2011) Probing the Interaction of the Diarylquinoline TMC207 with Its Target Mycobacterial ATP Synthase. PLoS ONE 6: e23575

Hartkoorn R, Sala, C., Neres, J., Pojer, F., Magnet, S., Mukherjee, R., Uplekar, S., Boy-Röttger, S., Altmann, KH., Cole, ST. (2012) Towards a new tuberculosis drug: pyridomycin – nature’s isoniazid. EMBO Mol Med 4: 1032–1042

Hu Y, Butcher, PD., Sole, K., Mitchison, DA., Coates, AR. (1998) Protein synthesis is shutdown in dormant Mycobacterium tuberculosis and is reversed by oxygen or heat shock. FEMS Microbiol Lett 158: 139145

Iacobino A, Piccaro, G., Giannoni, F., Mustazzolu, A., Fattorini, L. (2017) Mycobacterium tuberculosis Is Selectively Killed by Rifampin and Rifapentine in Hypoxia at Neutral pH. Antimicrob Agents Chemother 61: e02296–02316

Iacobino A, Piccaro, G., Giannoni, F., Mustazzolu, A., Fattorini, L. (2016) Activity of drugs against dormant Mycobacterium tuberculosis. Int J Mycobacteriol 5: S94–S95

Ignatov D, Salina, EG., Fursov, MV., Skvortsov, TA., Azhikina, TL., Kaprelyants, AS. (2015) Dormant non-culturable Mycobacterium tuberculosis retains stable low-abundant mRNA. BMC Genomics 16: 954–967

Jackson M (2014) The Mycobacterial Cell Envelope—Lipids. Cold Spring Harb Perspect Med 4: a021105

Jackson M, Stadthagen, G., Gicquel, B. (2007) Long-chain multiple methyl-branched fatty acid-containing lipids of Mycobacterium tuberculosis: Biosynthesis, transport, regulation and biological activities. Tuberculosis 87: 78–86

Jakkala K, Ajitkumar, P. (2019) Hypoxic Non-replicating Persistent Mycobacterium tuberculosis Develops Thickened Outer Layer That Helps in Restricting Rifampicin Entry. Front Microbiol 10: 2339

Kelley L, Mezulis, S., Yates, CM., Wass, MN., Sternberg, MJE. (2015) The Phyre2 web portal for protein modeling, prediction and analysis. Nat Protocols 10: 845–858

Kendall S, Movahedzadeh, F., Rison, SC., Wernisch, L., Parish, T., Duncan, K., Betts, JC., Stoker, NG. (2004) The Mycobacterium tuberculosis dosRS two-component system is induced by multiple stresses. Tuberculosis (Edinb) 84: 247–255

Kester J, Fortune, SM. (2014) Persisters and beyond: mechanisms of phenotypic drug resistance and drug tolerance in bacteria. Crit Rev Biochem Mol Biol 49: 91–101

Köhler G, Milstein, C. (1975) Continuous cultures of fused cells secreting antibody of predefined specificity. Nature 256: 495–497

Koul A, Vranckx, K., Dhar, N., Göhlmann, HWH., Özdemir, E., Neefs, JM., Schulz, M., Lu, P., Mørtz, E., McKinney, JD., Andries, K., Bald, D. (2014) Delayed bactericidal response of Mycobacterium tuberculosis to bedaquiline involves remodelling of bacterial metabolism. Nat Comm 5

Laalami S, Grentzmann, G., Bremaud, L. & Cenatiempo, Y. (1996) Messenger RNA translation in prokaryotes: GTPase centers associated with translational factors. Biochimie 78: 577–589

Lakshminarayana S, Tan, BH., Ho, PC., Manjunatha, UH., Dartois, V., Dick, T., Rao, SPS. (2015) Comprehensive physicochemical, pharmacokinetic and activity profiling of anti-TB agents. Journal of Antimicrobial Chemotherapy 70: 857–867

Leipe D, Wolf, YI., Koonin, EV., Aravind, L. (2002) Classification and evolution of P-loop GTPases and related ATPases. J Mol Biol 317: 41–72

Leistikow R, Morton, RA., Bartek, IL., Frimpong, I., Wagner, K., Voskuil, MI. (2010) The Mycobacterium tuberculosis DosR Regulon Assists in Metabolic Homeostasis and Enables Rapid Recovery from Nonrespiring Dormancy. J Bacteriol 192: 1662–1670

Li X, Sun, Q., Jiang, C., Yang, K., Hung, LW., Zhang, J, Sacchettini, JC. (2015) Structure of Ribosomal Silencing Factor Bound to Mycobacterium tuberculosis Ribosome. Structure 23: 1858–1865

Lin W, Sessions, PFD., Teoh, GHK., Mohamed, ANN., Zhu, OY., Koh, VHQ., Lay, MTA., Dedon, PC., Hibberd, ML., Alonso, S. (2016) Transcriptional Profiling of Mycobacterium tuberculosis Exposed to In Vitro Lysosomal Stress. Infection and Immunity 84: 2505–2523

Liu S, Shah, SJ., Wilmes, LJ., Feiner, J., Kodibagkar, VD., Wendland, MF., Mason, RP., Hylton, N., Hopf, HW., Rollins, MD. (2011) Quantitative tissue oxygen measurement in multiple organs using 19F MRI in a rat model. Preclinical and Clinical Imaging 66: 1722–1730

Loebel R, Shorr, E., Richardson, HB. (1933) The Influence of Foodstuffs upon the Respiratory Metabolism and Growth of Human Tubercle Bacilli. J Bacteriol 26: 139–166

Manabe Y, Bishai, WR. (2006) Latent Mycobacterium tuberculosis-persistence, patience, and winning by waiting. Nat Med 6: 1327–1329

Manjunatha U, Boshoff, HI., Barry, CE. (2009) The mechanism of action of PA-824: Novel insights from transcriptional profiling. Commun Integr Biol 2: 215–218

Maxson S, Schutze, GE., Jacobs, RF. (1994) Mycobacterium abscessus osteomyelitis: treatment with clarithromycin. Infect Dis Clin Pract 3: 203–205

McCune R, Feldmann, FM., Lambert, HP., McDermott, W. (1966) Microbial persistence. I. The capacity of tubercle bacilli to survive sterilization in mouse tissues. J Exp Med 123: 445–468

McCune R, Tompsett, R. (1956) Fate of Mycobacterium tuberculosis in mouse tissues as determined by the microbial enumeration technique. I. The persistence of drug-susceptible tubercle bacilli in the tissues despite prolonged antimicrobial therapy. J Exp Med 104: 737–762

Miranda-CasoLuengo A, Staunton, PM., Dinan, AM., Lohan, AJ., Loftus, BJ. (2016) Functional characterization of the Mycobacterium abscessus genome coupled with condition specific transcriptomics reveals conserved molecular strategies for host adaptation and persistence. BMC Genomics 17: 553

Mishra S, Ahmed, T., Tyagi, A., Shi, J., Bhushan, S. (2018) Structures of Mycobacterium smegmatis 70S ribosomes in complex with HPF, tmRNA, and P-tRNA. Sci Rep 8: 13587

Morris R, Nguyen, L., Gatfield, J., Visconti, K., Nguyen, K., Schnappinger, D., Ehrt, S., Liu, Y., Heifets, L., Pieters, J., Schoolnik, G., Thompson, CJ. (2005) Ancestral antibiotic resistance in Mycobacterium tuberculosis. Proc Natl Acad Sci U S A 102: 12200–12205

Mushatt D, Witzig, RS. (1995) Successful treatment of Mycobacterium abscessus infections with multidrug regimens containing clarithromycin. Clin Infect Dis 20: 1441–1442

Nicolle D, Fremond, C., Pichon, X., Bouchot, A., Maillet, I., Ryffel, B., Quesniaux, VJF. (2004) Long-term control of Mycobacterium bovis BCG infection in the absence of Toll-like receptors (TLRs): investigation of TLR2-, TLR6-, or TLR2-TLR4-deficient mice. Infect Immun 72: 6994–7004

Nuermberger E, Yoshimatsu, T., Tyagi, S., O’Brien, RJ., Vernon, AN., Chaisson, RE., Bishai, WR., Grosset, JH. (2004) Moxifloxacin-containing Regimen Greatly Reduces Time to Culture Conversion in Murine Tuberculosis. Am J Respir Crit Care Med 69: 421–426

Oh E, Becker, AH., Sandikci, A., Huber, D., Chaba, R., Gloge, F., Nichols, RJ., Typas, A., Gross, CA., Kramer, G., Weissman, JS., Bukau, B. (2011) Selective ribosome profiling reveals the cotranslational chaperone action of trigger factor in vivo. Cell 147: 1295–1308

Orme I, Basaraba, RJ. (2014) The formation of the granuloma in tuberculosis infection. Semin Immunol 26: 601–609

Park H, Guinn, KM., Harrell, MI., Liao, RL., Voskuil, MI., Tompa, M., Schoolnik, GK., Sherman, DR. (2003) Rv3133c/dosR is a transcription factor that mediates the hypoxic response of Mycobacterium tuberculosis. Mol Microbiol 48: 833–843

Parrish N, Dick, JD., Bishai, WR.. (1998) Mechanisms of latency in Mycobacterium tuberculosis. Trends Microbiol 6: 107–112

Piccaro G, Poce, G., Biava, M., Giannoni, F., Fattorini, L. (2015) Activity of lipophilic and hydrophilic drugs against dormant and replicating Mycobacterium tuberculosis. J Antibiot (Tokyo) 68: 711–714

Rao S, Alonso, S., Rand, L., Dick, T., Pethe K. (2007) The protonmotive force is required for maintaining ATP homeostasis and viability of hypoxic, nonreplicating Mycobacterium tuberculosis. Proc Natl Acad Sci U S A 105 11945–11950

Roberts D, Liao, RLP., Wisedchaisri, G., Hol, WGJ., Sherman, DR. (2004) Two Sensor Kinases Contribute to the Hypoxic Response of Mycobacterium tuberculosis*. J Bio Chem 279: 3082–23087

Rudra P, Hurst-Hess, KR., Cotten, KL., Miranda, Ap., Ghosh, P. (2020) Mycobacterial HflX is a ribosome splitting factor that mediates antibiotic resistance. PNAS 117: 629–634

Russell D (2007) Who puts the tubercle in tuberculosis? Nat Rev Microbiol 5: 39–47

Rustad T, Sherrid, AM., Minch, KJ., Sherman, DR. (2009) Hypoxia: a window into Mycobacterium tuberculosis latency. Cell Microbiol 11: 1151–1159

Saini D, Malhotra, V., Dey, D., Pant, NH., Das, TK., Tyagi, JS. (2004) DevR–DevS is a bona fide two-component system of Mycobacterium tuberculosis that is hypoxia-responsive in the absence of the DNA-binding domain of DevR. Microbiol 150: 865–875

Sassetti C, Boyd, DH., Rubin, EJ. (2003) Genes Required for Mycobacterial Growth Defined by High Density Mutagenesis. Mol Microbiol 48: 77–84

Sawyer E, Grabowska, AD., Cortes, T. (2018) Translational regulation in mycobacteria and its implications for pathogenicity. Nucleic Acids Res 46: 6950–6961

Schnappinger D, Ehrt, S., Voskuil, MI., Liu, Y., Mangan, JA., Monahan, IM., Dolganov, G., Efron, B., Butcher, PD., Nathan, C., Schoolnik, GK. (2003) Transcriptional Adaptation of Mycobacterium tuberculosis within Macrophages: Insights into the Phagosomal Environment. J Exp Med 198: 693–704

Shamimuzzaman M, Vodkin, L. (2018) Ribosome profiling reveals changes in translational status of soybean transcripts during immature cotyledon development. PLoS One 13: e0194596

Sharma S, Tyagi, JS. (2016) Mycobacterium tuberculosis DevR/DosR Dormancy Regulator Activation Mechanism: Dispensability of Phosphorylation, Cooperativity and Essentiality of α10 Helix. PLoS One 11: e0160723

Sherman D, Voskuil, M., Schnappinger, D., Liao, R., Harrell, MI., Schoolnik, GK. (2001) Regulation of the Mycobacterium tuberculosis hypoxic response gene encoding alpha-crystallin Proc Natl Acad Sci U S A 98: 7534–7539

Sherrid A, Rustad, TR., Cangelosi, GA., Sherman, DR. (2010) Characterization of a Clp Protease Gene Regulator and the Reaeration Response in Mycobacterium tuberculosis. PLoS ONE 5: e11622

Shi L, Sohaskey, CD., Kana, BD., Dawes, S., North, RJ., Mizrahi, V., Gennaro, ML. (2005) Changes in energy metabolism of Mycobacterium tuberculosis in mouse lung and under in vitro conditions affecting aerobic respiration. Proc Natl Acad Sci U S A 102: 15629–15634

Shleeva M, Kudykina, YK., Vostroknutova, GN., Suzina, NE., Mulyukin, AL., Kaprelyants, AS. (2011) Dormant ovoid cells of Mycobacterium tuberculosis are formed in response to gradual external acidification. Tuberculosis (Edinb) 91: 146–154

Soding J (2005) Protein homology detection by HMM-HMM comparison. Bioinformatics 21: 951–960

Sousa E, Tuckerman, JR., Gonzalez, G., Gilles-Gonzalez, MA. (2007) DosT and DevS are oxygen-switched kinases in Mycobacterium tuberculosis. Protein Sci 16: 1708–1719

Sprang S (1997) G Protein Mechanisms: Insights From Structural Analysis. Annu Rev Biochem 66: 639–678

Starosta A, Lassak, J., Jung, K., Wilson, DN. (2014) The bacterial translation stress response. FEMS Microbiol Rev 38: 1172–1201

Stothard P (2000) The sequence manipulation suite: JavaScript programs for analyzing and formatting protein and DNA sequences. Biotechniques 28: 1102–1104

Stover C, de la Cruz, VF., Fuerst, TR., Burlein, JE., Benson, LA., Bennett, LT., Bansal, GP., Young, JF., Lee, MH., Hatfull, GF. (1991) New use of BCG for recombinant vaccines. Nature 351: 456–460

Tomasz A, Albino, A., Zanati, E. (1970) Multiple antibiotic resistance in a bacterium with suppressed autolytic system. Nature 227: 138–140

Trauner A, Lougheed, KEA., Bennett, MH., Hingley-Wilson, SM., Williams, HD. (2012) The Dormancy Regulator DosR Controls Ribosome Stability in Hypoxic Mycobacteria. J Bio Chem 287: 24053–24063

Vaara M (1992) Agents that increase the permeability of the outer membrane. Microbiol Rev 56: 395–411

Velayati A, Farnia, P., Masjedi, MR., Zhavnerko, GK., Merza, MA., Ghanavei, J. (2011) Sequential adaptation in latent tuberculosis bacilli: observation by atomic force microscopy (AFM). Int J Clin Exp Med 4: 193–199

Verstraeten N, Fauvart, M., Versées, W., Michiels, J. (2011) The Universally Conserved Prokaryotic GTPases. Microbiol Mol Biol Rev 75: 507–542

Wada A, Igarashi, K., Yoshimura, S., Aimoto, S., Ishihama, A. (1995) Ribosome modulation factor: stationary growth phase-specific inhibitor of ribosome functions from Escherichia coli. Biochem Biophys Res Commun 214: 410–417

Wayne L, Hayes, LG. (1996) An in vitro model for sequential study of shiftdown of Mycobacterium tuberculosis through two stages of nonreplicating persistence. Infect Immun 64: 2062–2069

Wayne L, Sohaskey, CD. (2001) Nonreplicating persistence of mycobacterium tuberculosis. Annu Rev Microbiol 55: 139–163

Yamagishi M, Matsushima, H., Wada, A., Sakagami, M., Fujita, N., Ishihama, A. (1993) Regulation of the Escherichia coli rmf gene encoding the ribosome modulation factor: growth phase- and growth rate-dependent control. EMBO J 12: 625–630

Yokoyama W, Christensen M, Dos Santos G., Miller, D., Ho, J., Wu, T., Dziegelewski, M., Neethling, FA. (2013) Production of monoclonal antibodies. Curr Protoc Immunol 102

Zhang Y, Mandava, C. S., Cao, W., Li, X., Zhang, D., Li, N., Zhang, Y., Zhang, X., Qin, Y., Mi, K., Lei, J., Sanyal, S. & Gao, N. (2015) HflX is a ribosome-splitting factor rescuing stalled ribosomes under stress conditions. Nat Struct Mol Biol 22: 906–913

